# Packed for Ossification: High-Density Bioprinting of hPDC Spheroids in HAMA for Endochondral Ossification

**DOI:** 10.1101/2025.09.09.674866

**Authors:** Ane Albillos Sanchez, Maria Paula Marks, Paula Casademunt, Adrián Seijas-Gamardo, Ioannis Papantoniou, Lorenzo Moroni, Carlos Mota

**Affiliations:** Complex Tissue Regeneration Department, MERLN Institute for Technology-Inspired Regenerative Medicine, Maastricht University, 6229 ER, Maastricht, The Netherlands; Department of Cell Biology-Inspired Tissue Engineering, MERLN Institute for Technology-Inspired Regenerative Medicine, Maastricht University, 6229 ER, Maastricht, The Netherlands; Universitat Pompeu Fabra, Physense, BCN Medtech, Department of Engineering, 08018, Barcelona, Spain; Prometheus Division of Skeletal Tissue Engineering, KU Leuven, O&N1, Herestraat 49, PB 813, 3000 Leuven, Belgium; Department of Development and Regeneration, Skeletal Biology and Engineering Research Center, KU Leuven, O&N1, Herestraat 49, PB 813, 3000 Leuven, Belgium; Institute of Chemical Engineering Sciences, Foundation for Research and Technology-Hellas, Stadiou 26504, Platani, Patras, Greece

**Keywords:** Human Periosteum-Derived Stem Cells (hPDCs), Hyaluronic Acid-Based Hydrogels, Endochondral Ossification, Extrusion-Based Bioprinting, Spheroids

## Abstract

Long bone fractures are primarily repaired through endochondral ossification, a process in which a soft cartilage template forms at the injury site and is gradually replaced by bone. While bone has an innate self-healing capacity, this process can be disrupted in cases of large or complex defects, where regeneration fails, and clinical intervention is required. This study aimed at the development of a tissue engineering approach using human periosteum-derived cell (hPDC) spheroids encapsulated or bioprinted at high density within hyaluronic acid methacrylate (HAMA) hydrogels to support hypertrophic cartilage formation as a template for endochondral bone regeneration. We first compared different encapsulation time points (days 1, 7, and 14), finding that early encapsulation (day 1) enhanced spheroid fusion, increased DNA content, and promoted hypertrophic cartilage formation, as indicated by greater glycosaminoglycan (GAG) and collagen deposition along with *lacunae* formation. Next, HAMA-encapsulated spheroids were compared to spheroids formed using a standardized microwell platform, demonstrating that encapsulation promoted a more mature cartilage-like matrix with thicker collagen fibers and enhanced hypertrophic differentiation. Gene expression and immunostaining confirmed progression toward hypertrophic and osteogenic phenotypes. Finally, extrusion-based bioprinting of HAMA bioinks comprising a high-density of hPDC spheroids demonstrated scalability, improved spheroid alignment, and maintained robust cell viability and hypertrophic differentiation. HA’s bioactivity and regulatory advantages support clinical translation, although achieving spatial control remains an area for further optimization.

**Graphical Abstract:** 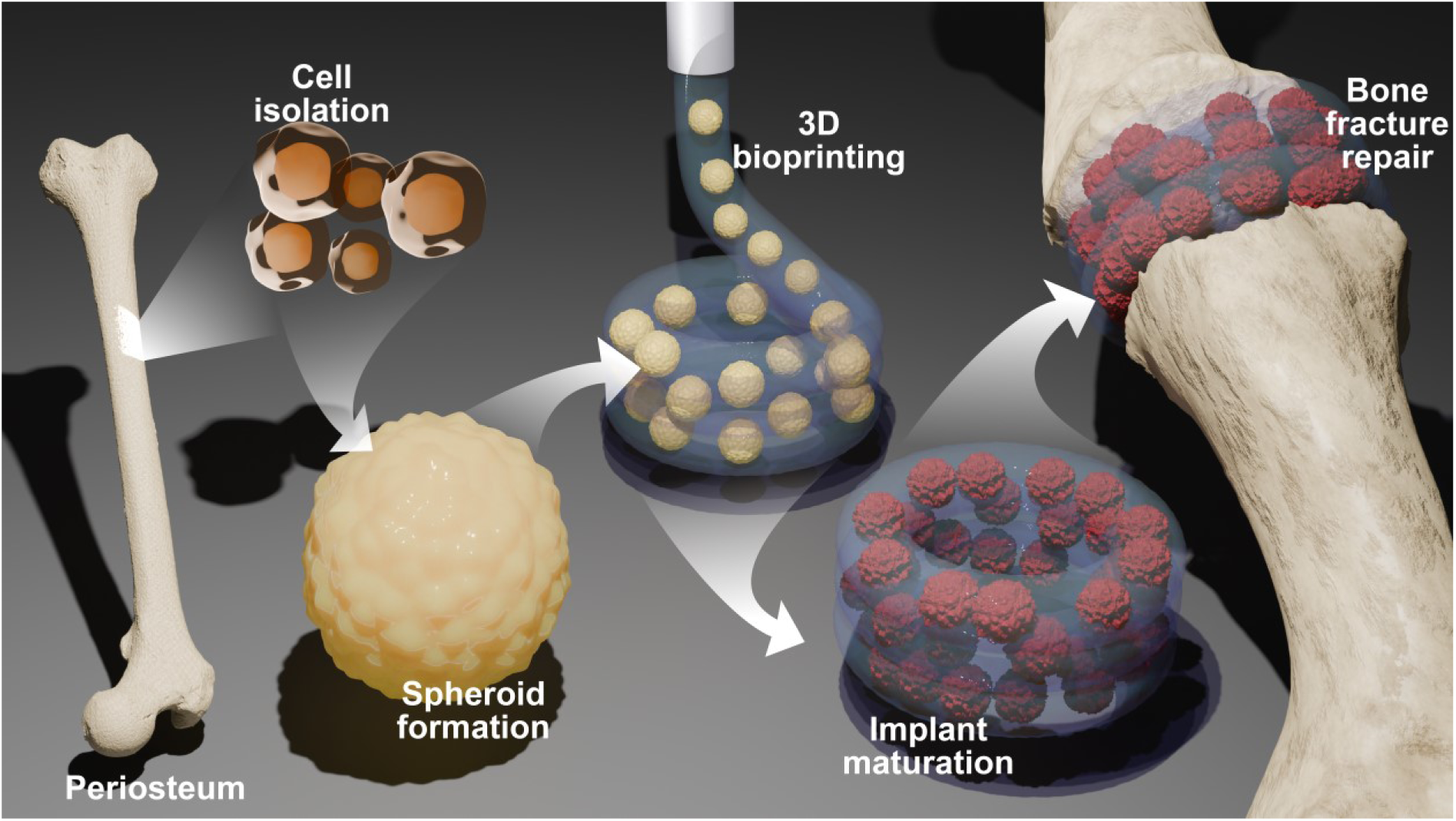

## 1. Introduction

Bone, in contrast to many other tissues, has a self-repairing capacity for non-critical size defects, healing without scarring and restoring its previous structure and strength [1, 2]. Both intramembranous and endochondral ossification pathways play important roles in this precisely regulated fracture healing process, which follows routes like those that take place during fetal bone development [3]. Endochondral ossification plays a particularly important role in the repair of long bone fractures. A soft cartilage callus develops at the fracture site and is gradually replaced by mineralized bone [4]. However, this process is limited to smaller injuries; critical size defects exceed the bone’s intrinsic regenerative capacity and do not spontaneously heal through endochondral ossification. As a result, complications such as delayed union or non-union still occur in up to 13.3% of tibial fractures [5], especially under conditions such as acute trauma, infection, or altered biological environments [6]. Autograft, allograft, and synthetic implants are examples of traditional therapies for impaired fracture healing; nevertheless, they have drawbacks, including donor-site morbidity, immunological rejection, and poor integration [7]. This highlights the urgent need for improved alternative methods to promote long bone repair. Tissue engineering and biofabrication aim to produce functional bone grafts that replicate the natural structure and mechanical characteristics of the bone tissue, becoming potential solutions for addressing these limitations [8–10]. To guide tissue growth and regeneration, developmental engineering integrates concepts from developmental biology. It seeks to replicate the formation of a cartilaginous soft callus that progressively transforms into bone during the process of endochondral ossification [11, 12]. For bone formation to be successful, this process needs to include phases of cellular condensation, self-assembly, and matrix maturation [13]. Reproducing this complex spatiotemporal pattern *in vitro* has great potential for regenerative medicine, even though it remains highly challenging due to the need for precise coordination of cellular response within a controlled 3D environment [11, 12, 14].

Mesenchymal stem cells present in the periosteum, the thin vascularized membrane covering the surfaces of bones, are known as human periosteum-derived stem cells or hPDCs. HPDCs can differentiate into osteogenic, chondrogenic, and adipogenic lineages [15, 16]. Importantly, as hPDCs are necessary for the natural processes of bone growth and repair, they are highly relevant for developmental engineering approaches aimed at bone regeneration [17]. Using hPDCs in spheroid form has demonstrated great promise for developmental engineering. This approach mimics the natural development of a cartilaginous callus, which is crucial for bone repair, by using cell aggregation and condensation, two crucial early stages in endochondral ossification [18–21].

Spheroids have been investigated in the context of musculoskeletal tissue engineering and offer substantial benefits when incorporated into biofabrication and bioprinting techniques [11]. Since they may be deposited layer by layer in exact spatial configurations, spheroids serve as tissue building blocks that can fuse into larger, more organized structures that aid in the processes of growth and maturation [22, 23]. For this application, extrusion-based bioprinting offers a particularly versatile method as it makes use of spheroid-based bioinks that can be extruded through a nozzle to create continuous, cell-rich structures. In addition to being effective and scalable, this technique makes tissue construct production easier by enabling individual spheroid fusion on a receiving substrate [24].

Hyaluronic acid (HA), particularly in its sulfated form, plays a crucial role in bone regeneration by influencing cell activity and activating signaling pathways such as TGFβ, BMP, and Wnt, which are essential for chondrogenesis and endochondral ossification [25, 26]. These pathways promote chondrocyte differentiation and matrix maturation, key steps in forming a cartilage template for bone development. The benefits of utilizing HA hydrogels in combination with human bone marrow-derived mesenchymal stem cells (hBMSCs) to promote hypertrophic cartilage production *in vitro*, which can subsequently go through endochondral ossification *in vivo*, are highlighted by Yamazaki *et al.* [27]. A scalable method for producing integrated, vascularized bone tissue is offered by the HA hydrogel’s capacity to promote homogeneous cell distribution and regulated differentiation, which has great promise for clinical uses in bone regeneration and repair [27].

This study aimed to explore the use of hPDC spheroids combined with HA methacrylate (HAMA), a photo-crosslinkable form of HA, to promote hypertrophic cartilage formation *in vitro*. We investigated whether early-stage encapsulation in HAMA could enhance key processes in endochondral ossification, such as spheroid fusion, condensation, and hypertrophic differentiation, compared to standard microwell culture. Building on these findings, the approach was translated into a high-density bioprinting platform to assess its potential for scalable and clinically relevant bone tissue engineering applications.

## 2. Materials and Methods

### 2.1. Cell culture and expansion

Human periosteum-derived cells (hPDCs) were kindly provided by Prometheus, the Skeletal Tissue Engineering division of Katholieke Universiteit Leuven. hPDCs cell pool was obtained from five donors (14 ± 3 years old) and isolated from periosteal tissues as previously described [28]. The cells were expanded from passage 6 to passage 8 under standard culture conditions. For expansion, a high-glucose Dulbecco’s Modified Eagle’s Medium (DMEM) with pyruvate (11995065, Gibco) was used, supplemented with 10% fetal bovine serum (FBS, F7524, Sigma-Aldrich) and 1% antibiotic/antimycotic solution (15240062, Gibco). Culturing was carried out under standard conditions in a humidified incubator at 37°C with 5% CO₂, with medium changes performed every three to four days, until 90% confluency was reached. At this point, cells were harvested using TrypLE Express (12605010, Gibco).

All procedures were approved by the Ethical Committee for Human Medical Research (KU Leuven) and informed consent was obtained from all patients (ML7861).

### 2.2. Spheroid generation and differentiation in chondrogenic medium

To generate hPDC spheroids, cells were harvested and pelleted by centrifugation at 300 × g for 5 minutes. The cell pellet was then resuspended in fresh medium at a concentration of 1.5 × 10⁵ cells/mL. A 2 mL aliquot of the resulting cell suspension was seeded into each well of commercially available standardized microwell plates containing 1200 microwells per well (AggreWell 400, 34415, STEMCELL Technologies), ensuring an average of 250 cells per spheroid.

Following seeding, the cells were incubated at 37°C to promote cell sedimentation and spheroid formation. Spheroid differentiation was evaluated over 21 days of culture in a hypertrophic chondrogenic differentiation medium (HCM). The HCM consisted of LG-DMEM (11885084, Gibco) supplemented with 1% antibiotic-antimycotic (100 units/mL penicillin, 100 mg/mL streptomycin, and 0.25 mg/mL amphotericin B, 15240062, Gibco), 100 nM dexamethasone (D4902, Merck), 1 mM ascorbate-2 phosphate (A8960, Merck), 40 µg/mL proline (P5607, Merck), ITS+ Premix Universal Culture Supplement (354352, Corning) (containing 6.25 µg/mL insulin, 6.25 µg/mL transferrin, 6.25 µg/mL selenious acid, 1.25 µg/mL bovine serum albumin (BSA), and 5.35 µg/mL linoleic acid), 20 µM Rho-kinase inhibitor Y27632 (1683, Axon Medchem), 100 ng/mL GDF5 (120-01, PeproTech), 100 ng/mL BMP-2 (INDUCTOS^®^), 10 ng/mL TGFβ1 (100-21, PeproTech), 1 ng/mL BMP-6 (120-06, PeproTech), and 0.2 ng/mL FGF-2 (100-18C, PeproTech).

The culture medium was partially refreshed every three to four days, with half of the volume being replaced. Spheroids were harvested on days 1, 7, 14, and 21 during differentiation for further analysis.

### 2.3. Synthesis and characterization of methacrylated hyaluronic acid (HAMA)

The HAMA used in this study is from the same synthesized batch previously described in [29], ensuring consistency in material properties. The degree of methacrylation and baseline photo-rheological properties were reported in that study.

To identify suitable conditions for partial crosslinking before bioprinting, additional rheological measurements were performed. A solution containing 2% w/v HAMA and 0.01% w/v lithium phenyl-2,4,6-trimethylbenzoylphosphinate (LAP, 900889, Merck) in phosphate buffered saline (PBS, D8537, Sigma-Aldrich) was prepared and allowed to dissolve overnight at room temperature. Rheological analysis was conducted using a DHR-2 Rheometer (TA Instruments) equipped with a 20 mm cone plate geometry and a quartz bottom plate. UV light at 365 nm was delivered through a liquid light guide connected to an M365LP1 Mid Power Mounted LED UV-light attachment (Thorlabs). The LED intensity at the sample interface was regulated by a DC2200 1 Channel LED Driver (10A).

To ensure accurate intensity readings, calibration was performed using a fitted sensor from TA Instruments, specifically designed to align with the lower geometry and capture the actual intensity received by the sample. Measurements were conducted over a 5-minute time sweep at 22°C (measured room temperature), with a strain of 1% and a frequency of 1 Hz. Prior to initiating measurements, samples were stabilized for 1 minute. The UV source was then activated, emitting 365 nm light at an intensity of 12 mW/cm², and samples were exposed for durations of 30, 40, 50, and 60 seconds.

### 2.4. Encapsulation and differentiation of hPDC spheroids in HAMA

For the encapsulation experiments, hPDC spheroids were initially generated using microwell plates as previously described and cultured in HCM for 24 hours. Following this incubation period, spheroids were harvested and suspended in a 2% w/v HAMA and 0.1% w/v LAP solution in PBS at a density of 15,000 spheroids/mL.

Hydrogels were formed by dispensing 20 µL of the spheroid-laden HAMA mixture into a custom-made polydimethylsiloxane (PDMS) mold, with cavities having dimensions of 1 mm in height and 8 mm in diameter. The hydrogels were crosslinked using a Thorlabs’ DC4104 four-channel LED driver with UV light emitting at 365 nm with an intensity of 12 mW/cm^2^ for 30 seconds. Crosslinked spheroid-laden hydrogels were maintained in HCM for an additional period of 20 days. The medium was refreshed every three or four days. Samples were collected on days 1, 7, 14, and 21 to evaluate differentiation.

### 2.5. Bioprinting and differentiation of hPDC spheroids in HAMA

As previously described, hPDC spheroids were prepared using microwell plates and cultured in HCM for 24 hours. A HAMA solution was prepared at a concentration exceeding 2% w/v in PBS to account for subsequent dilution. This solution, containing 0.01% w/v LAP in PBS, was partially crosslinked in a 2 mL Eppendorf tube under 12 mW/cm² light exposure for 40 seconds. Following partial crosslinking, additional LAP was added to achieve a final concentration of 0.1% w/v in PBS, ensuring a final HAMA concentration of 2% w/v in PBS. The final solution was then mixed with spheroids at a density of 40,000 spheroids/mL.

A pressure-assisted extrusion bioprinter (BioScaffolder 3.1, Gesim - Gesellschaft für Silizium-Mikrosysteme mbH, Germany) was used to produce all constructs. The spheroid-laden HAMA solution was loaded into a 1 mL syringe at room temperature. This solution was extruded through a smooth flow tapered tip nozzle with an internal diameter of 410 µm (7005007, Nordson) at a controlled pressure of 10 kPa and a constant speed of 25 mm/s. The printing process was conducted at room temperature. After the extrusion process, the constructs were subjected to 30 s of UV crosslinking (365 nm, 12 mW/cm^2^).

The bioprinted constructs were 4 mm in radius and consisted of four layers, with a subsequent set of connected circles with an infill distance of 1 mm as a deposition pattern. Crosslinked bioprinted constructs were cultured in HCM for an additional 20 days, after which samples were collected for differentiation analysis (day 21). The medium was refreshed every three or four days.

### 2.6. Viability assay

The viability of hPDC spheroids was assessed at different time points under material-free conditions, as well as following encapsulation or bioprinting in HAMA. Samples were incubated with 2 µM Calcein-AM (C3099, Invitrogen) for 45 minutes at 37°C, and 4 µM Ethidium Homodimer-1 (EthD-1, E1169, Invitrogen) was added during the last 15 minutes of incubation.

Fluorescence imaging was performed using an automated inverted Nikon Ti-E microscope, equipped with a Lumencor Spectra X light source, a Photometrics Prime 95B sCMOS camera, an MCL NANO Z500-N TI z-stage, and an Okolab incubator (37°C, 5% CO2) to facilitate live cell imaging.

### 2.7. DMMB assay and DNA quantification

The amount of sulfated glycosaminoglycans (GAGs) secreted into the culture media by microwell-cultured or HAMA-encapsulated spheroids was quantified using the dimethyl-methylene blue (DMMB) assay (pH 1.5) [30]. For microwell-cultured samples, the spheroids from one well (∼ 1200), were pooled to represent one replicate. Absorbances were measured at 525 and 595 nm using a CLARIOstar Plate Reader (BMG Labtech), and concentrations were calculated by comparing the ratio of absorbances to a linear standard curve prepared with chondroitin sulfate C (C4384, Sigma-Aldrich). To quantify the retained GAGs and DNA, samples were digested overnight in 1 mg/mL proteinase K (P6556, Sigma-Aldrich) solution in Tris/EDTA buffer at 56°C. GAGs concentrations in the digested samples (retained GAGs) were measured using the same protocol as for the corresponding culture media supernatants from the same samples. For both secreted and retained GAGs, the amounts from spheroid-free gels were subtracted from the spheroid-laden constructs to determine the amount of GAGs produced by the cells.

For analyzing spheroids encapsulated in HAMA, the constructs were freeze-dried and pre-treated with overnight digestion in 1 mg/mL hyaluronidase type II (H2126, Sigma-Aldrich) at 37°C to facilitate cell release before conducting the assay. One hydrogel containing ∼ 300 spheroids represented one replicate.

GAGs content was normalized to DNA content, which was determined using the CyQUANT Cell Proliferation Assay Kit (C7026, Invitrogen). To degrade any residual RNA, the lysates were incubated for 1 hour at room temperature with a buffer containing RNase A (1:500 dilution) in 20X diluted cell lysis buffer. DNA content was then evaluated by incubating the samples with the kit’s fluorescent dye at a 1:1 ratio for 15 minutes, followed by fluorescence measurement (excitation/emission = 480/520 nm) using the same CLARIOstar Plate Reader. DNA concentrations were determined from a standard curve prepared using a 100 µg/mL sample of bacteriophage λ DNA provided in the kit.

### 2.8. Histology

For histological analysis, encapsulated/bioprinted and biomaterial-free spheroids were fixed using 3.6% v/v formaldehyde solution (252549, Sigma-Aldrich) in PBS for 30 minutes at room temperature. Following fixation, samples were washed with PBS, embedded in Epredia HistoGel Specimen Processing Gel (12006679, Thermo Fisher Scientific), and processed for paraffin embedding. Dehydration was performed through a graded ethanol series (50%, 70%, 96%, and 100%) followed by xylene (0215869204, VWR) treatment, before embedding in paraffin (1.116092504, VWR). Samples were sectioned with 6 μm thickness using an Epredia HM 355S Automatic Microtome (Thermo Fisher Scientific) and collected on glass slides (631-9483, VWR) for staining.

For staining, sections were deparaffinized in xylene and rehydrated through a graded series of ethanol solutions of decreasing concentrations (100%, 96%, 70%, 50%) and deionized water. For Haematoxylin and Eosin staining, the sections were immersed in Harris Hematoxylin (3801562, Leica Biosystems) for 4 minutes, followed by washing in running tap water for 10 minutes. The sections were then briefly dipped in acid alcohol and immersed in eosin Y solution (HT110232, Sigma-Aldrich) for 2 minutes. For Alcian Blue staining, the samples were immersed for 5 minutes in a 1% (w/v) Alcian Blue 8GX solution (A5268, Sigma-Aldrich) prepared in 0.1 M hydrochloric acid (pH 1), followed by washing in deionized water. Nuclei were counterstained with 0.1% (w/v) Nuclear Fast Red solution (N3020, Sigma-Aldrich) prepared in 5% (w/v) aluminium sulfate (A0843, Sigma-Aldrich) and applied for 5 minutes, followed by rinsing in deionized water. For Picro-Sirius Red staining, the samples were stained according to the manufacturer’s protocol (ab150681, Abcam). Briefly, sections were incubated in Picro-Sirius Red solution for 60 minutes, then rinsed with acetic acid solution followed by absolute ethanol. For Safranin O staining, the samples were stained in Weigert’s iron Hematoxylin (HT1079, Sigma) for 10 minutes. The samples were then washed in running tap water for 10 minutes and stained in 0.04% aqueous Fast Green solution (F7252, Sigma). The samples were rinsed in 1% acetic acid solution for 10 seconds and stained in 0.2% Safranin O solution (HT90432, Sigma) for 5 minutes.

After staining, the sections were dehydrated, immersed in xylene and mounted with a coverslip using dibutylphthalate polystyrene xylene (DPX, 06522, Sigma-Aldrich). The stained sections were imaged using an automated inverted Nikon Ti-E microscope, equipped with a Lumencor Spectra light source, a Nikon DS-Ri2 camera, and an MCL NANO Z200-N TI z-stage.

### 2.9. Immunofluorescence staining

The immunofluorescence staining protocol varied depending on the antibody type. For the nuclear markers Osterix, Runx2, and SOX-9, a full-mount staining was performed, while 6 μm sections were used for the extracellular matrix (ECM) markers Aggrecan, type X collagen and type I collagen. For sections, samples underwent the same fixation, dehydration, embedding, and sectioning protocols as previously mentioned. Full-mount samples were fixed using 3.6% v/v formaldehyde solution for 30 minutes at room temperature. All samples were permeabilized with 1% Triton X-100 solution (T8787, Merck) for 10 minutes at room temperature. Blocking was performed in PBS containing 1% bovine serum albumin (BSA, 421501J, Avantor), 5% goat serum (G9023, Sigma-Aldrich), and 0.1% Tween-20 (P9416, Merck). Full-mount samples were blocked for 24 hours at 4 °C, while sections were blocked for 1 hour at room temperature.

Samples were then incubated with primary antibodies at a 1:100 dilution for 24 hours at 4°C. These included rabbit anti-type I collagen (600-401-103-0.1, Rockland), rabbit anti-Runx2 (PA5-82787, Abcam), mouse anti-Osterix (MAB7547, R&D Systems), rabbit anti-SOX-9 conjugated to Alexa Fluor 647 (ab196184, Abcam), mouse anti-Collagen X (ab49945, Abcam) and mouse anti-Aggrecan (MA3-16888, Thermo Fisher Scientific). A rabbit IgG isotype control antibody (ab172730, Abcam) was also used. After washing, samples were incubated with secondary antibodies, goat anti-rabbit IgG H&L Alexa Fluor 647 (ab150079, Abcam) or goat anti-mouse IgG H&L Alexa Fluor 647 (ab150115, Abcam), at a 1:1000 dilution. For sections, incubation was performed for 2 hours at room temperature, while for full-mount samples, secondary antibody incubation was carried out for 24 hours at 4 °C.

Nuclei were counterstained with 4′,6-diamidino-2-phenylindole (DAPI, 32670, Merck) for 20 minutes at room temperature at a 1:200 dilution, while F-actin was stained with Alexa Fluor 488 Phalloidin (10125092, Thermo Fisher Scientific) at a 1:200 dilution for 1 hour at room temperature. After staining, full-mount samples were stored in PBS until imaging, whereas sectioned samples were mounted with coverslips using ProLong Gold Antifade Mountant (11559306, Invitrogen).

Imaging of sectioned samples was performed using an automated inverted Nikon Ti-E microscope equipped with a Lumencor Spectra X light source, a Photometrics Prime 95B sCMOS camera, and an MCL NANO Z500-N TI z-stage. Full-mount samples were imaged using a Tandem confocal system (Leica TCS SP8 STED) equipped with a white light laser (WLL). For full-mount samples, a Z-stack maximum projection was used to capture the entire depth of the sample. Excitation was carried out using dye-specific wavelengths from the WLL or a 405 nm photodiode for DAPI. Emissions were detected using photomultiplier tubes (PMTs) for DAPI or Hybrid Detectors (HyDs) for the 647 nm signal.

### 2.10. Image-Based Morphometric and Clustering Analysis of Spheroids

Phalloidin-stained confocal Z-stacks of hPDC spheroids were acquired at days 1, 7, 14, and 21 for both microwell-cultured and HAMA-encapsulated conditions. Maximum projection images were used for all analyses. For each condition and time point, six independent images were analyzed. Depending on spheroid density within each field of view, this yielded between 6 and 12 individual spheroids per condition. In cases where two or more spheroids were fused, they were treated as a single unit for analysis.

An automated image analysis pipeline was implemented in ImageJ to segment spheroids using Otsu’s thresholding. Morphological operators were applied to fill internal holes, and shape descriptors including area, roundness, and solidity were extracted from the resulting binary masks.

In parallel, deep image features were extracted from the same projections using a pretrained ResNet50 convolutional neural network. These features were used to construct a latent feature space via principal component analysis (PCA), enabling visualization of the distribution of spheroids based on high-dimensional image characteristics. K-means clustering (k = 3, selected using the elbow method) was applied to group morphologically similar spheroids. Representative examples from extreme and intermediate positions along PC1 and PC2 were selected to illustrate shape and actin organization.

### 2.11. PCR

Quantitative real-time polymerase chain reaction (qRT-PCR) was used to quantify mRNA of relevant markers for endochondral ossification. For the non-encapsulated condition, spheroids were pooled from 2 wells (∼ 2400 spheroids) to represent one replicate. Spheroids were washed in PBS followed by treatment with TRIzol reagent (15596026, Thermo Fisher Scientific). The total RNA was extracted using chloroform/isopropanol methodology. The RNA was reverse transcribed according to manufacturer protocol using iScript cDNA Synthesis Kit (1708890, Bio-Rad).

For HAMA hydrogel samples, 2 hydrogels were pooled, freeze-dried, and digested overnight in 1 mg/mL hyaluronidase type II (H2126, Sigma-Aldrich) at 37°C and 500 rpm using a thermomixer (ThermoMixer C, Eppendorf). After digestion and to facilitate RNA extraction, hydrogels were smashed using pellet pestles (11815125, Thermo Fisher Scientific) in a portion of TRIzol reagent before adding the whole volume. After extracting RNA as previously described, a SuperScript IV Single Cell/Low Input cDNA PreAmp Kit (11752048, Thermo Fisher Scientific) was used for cDNA synthesis and amplification from a low amount of total RNA. Finally, a GeneJET PCR Purification Kit (10400450, Thermo Fisher Scientific) was used to purify the obtained cDNA.

qRT-PCR was carried out using the Biorad-CFX96 detection system (3600037, Bio-Rad) with iQ SYBR Green Supermix (1708886, Bio-Rad). Gene expression was normalized to the geometric mean of housekeeping genes Receptor Accessory Protein 5 (*REEP5*) and Proteasome Subunit Beta type-2 (*PSMB2)*, with relative differences calculated using the 2−ΔΔCt method, with expression levels normalized to day 1. For *COL2A1* in the microwell condition, data was compared to day 7 because transcripts were not detected on day 1. The primer sequences used are reported in Table S1.

### 2.12. Statistical analysis

Statistical analyses were performed using GraphPad Prism (version 10.3.1). An unpaired t-test was applied to compare differences between two groups, while one-way ANOVA followed by Tukey’s post-hoc test was used for comparisons among three or more groups. Statistical significance was defined as a p-value below 0.05. Data are presented as bar graphs showing the mean ± SEM, with each graph based on three independent replicates (N = 3). Statistical significance is denoted by asterisks: (*) p < 0.05, (**) p < 0.01, (***) p < 0.001, and (****) p < 0.0001, providing a visual indication of the significance levels.

## 3. Results

### 3.1. Optimization of Encapsulation Timepoint for Spheroid Differentiation and Fusion Capacity within HAMA

hPDC spheroids were encapsulated at different stages of differentiation to evaluate the optimal time point to promote both differentiation and fusion. These stages corresponded to the duration of spheroid culture in hypertrophic chondrogenic medium (HCM) within microwell plates prior to encapsulation. Microtissues were encapsulated in HAMA on days 1, 7, and 14, followed by continued culture in HCM until day 21 (schematically depicted in Figure 1A).

**Figure 1.**
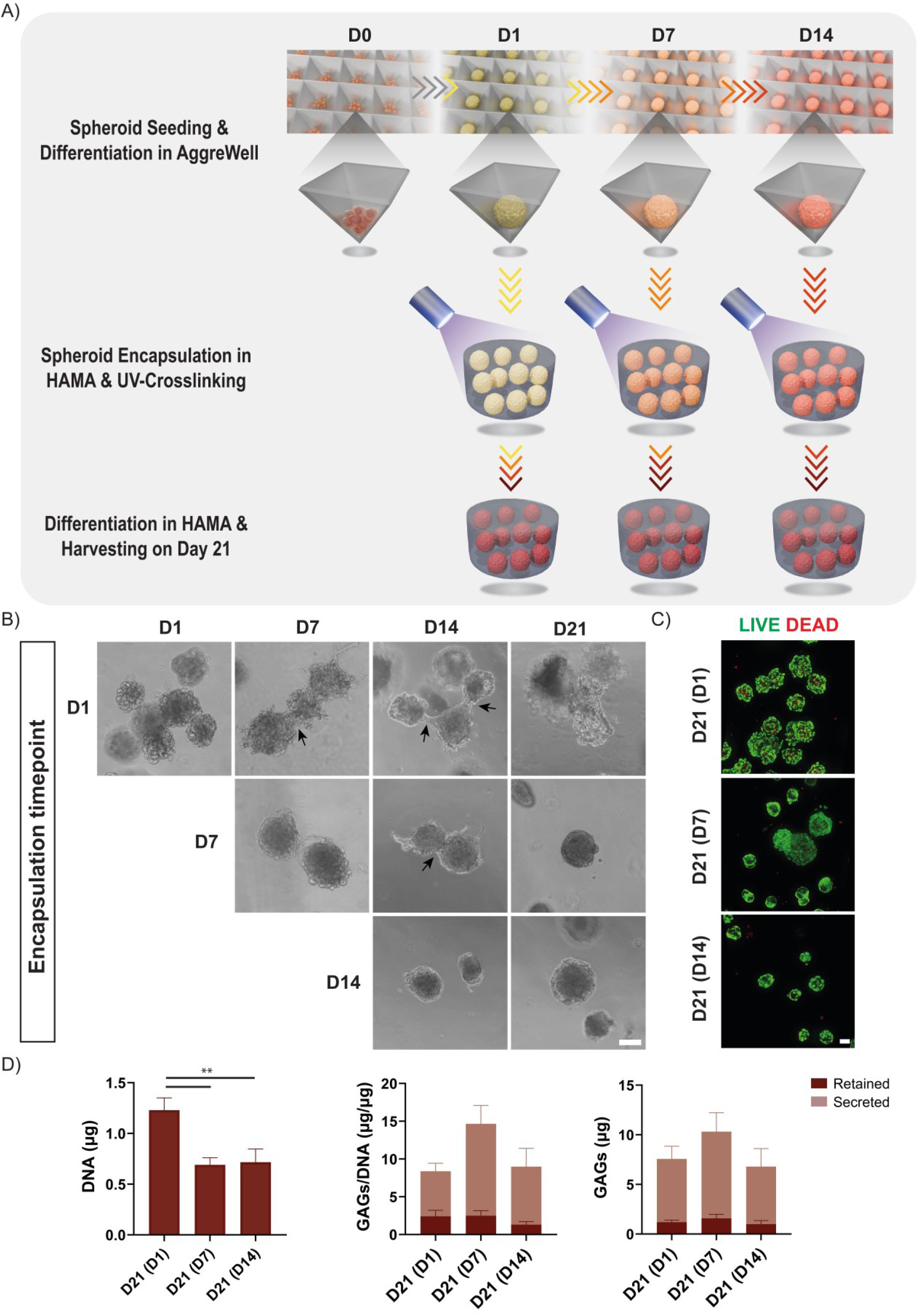
(A) Overview of experimental timeline. hPDC spheroids were formed and differentiated in microwell plates and encapsulated in HAMA hydrogels at day 1, 7, or 14. All groups were cultured until day 21 to assess fusion and hypertrophic differentiation. (B) Bright-field images of spheroids encapsulated at different time points, taken every 7 days, showing their structural organization and fusion over time. Examples of individual spheroids merging together are highlighted with black arrows. (C) Live-dead staining representative images showing cell viability at day 21 for spheroids encapsulated on day 1 (D21(D1)), day 7 (D21(D7)) and day 14 (D21(D14)). Live cells were stained with Calcein-AM (green), and dead cells were stained with Ethidium Homodimer-1 (EthD-1, red). (D) Quantification of DNA content (left), sulfated glycosaminoglycans (GAGs) normalized to DNA (middle), and total GAGs (right) at day 21 for spheroids encapsulated on day 1 (D21(D1)), day 7 (D21(D7)) and day 14 (D21(D14)). The GAG/DNA graph displays stacked bars representing both retained and secreted levels. Data are presented as mean ± SEM (*N* = 3). Statistical significance was determined using one-way ANOVA followed by Tukey’s post-hoc multiple comparison test (\*\**p* < 0.01). Scale bars represent 100 µm.

Spheroids encapsulated on day 1 showed a greater degree of fusion compared to those encapsulated on day 7, as evidenced by the merging of individual spheroids, while spheroids encapsulated on day 14 showed no fusion capacity (Figure 1B). Live-dead staining on day 21 showed high cell viability across all three encapsulation time points, indicating that the encapsulation process and the HAMA hydrogel environment supported cell survival in all tested conditions (Figure 1C).

Quantification of DNA content revealed that spheroids encapsulated on day 1 contained significantly higher DNA amounts by day 21 when compared with those encapsulated on day 7 or day 14 (Figure 1D, left). Despite differences in DNA content, GAGs content normalized to DNA (Figure 1D, middle) and total GAGs content (Figure 1D, right) were not significantly different across the three encapsulation timepoints. This lack of significant difference was observed both for GAGs retained within the constructs and those secreted into the surrounding medium. This indicates that the timepoint of encapsulation did not significantly affect the extracellular matrix (ECM) secretion of the encapsulated spheroids.

The ECM composition and tissue organization of hPDC spheroids encapsulated at different time points were analyzed through histological staining at day 21 using Hematoxylin and Eosin (HE), Alcian Blue (AB), Safranin O (SAF O), and Picro-Sirius Red (PR), with PR also visualized under polarized light ((PR)PL) (Figure 2A). GAG deposition was observed in blue with AB staining, both around and within the spheroids. However, SAF O staining, which specifically binds to sulfated proteoglycans, was not clearly detected for all conditions. PR staining, which detects collagen, was more intense in spheroids encapsulated on day 1, particularly when viewed under polarized light. *Lacunae*, a hallmark of cartilage maturation, were present in spheroids encapsulated on day 1 (Figure 2A, PR staining).

**Figure 2.**
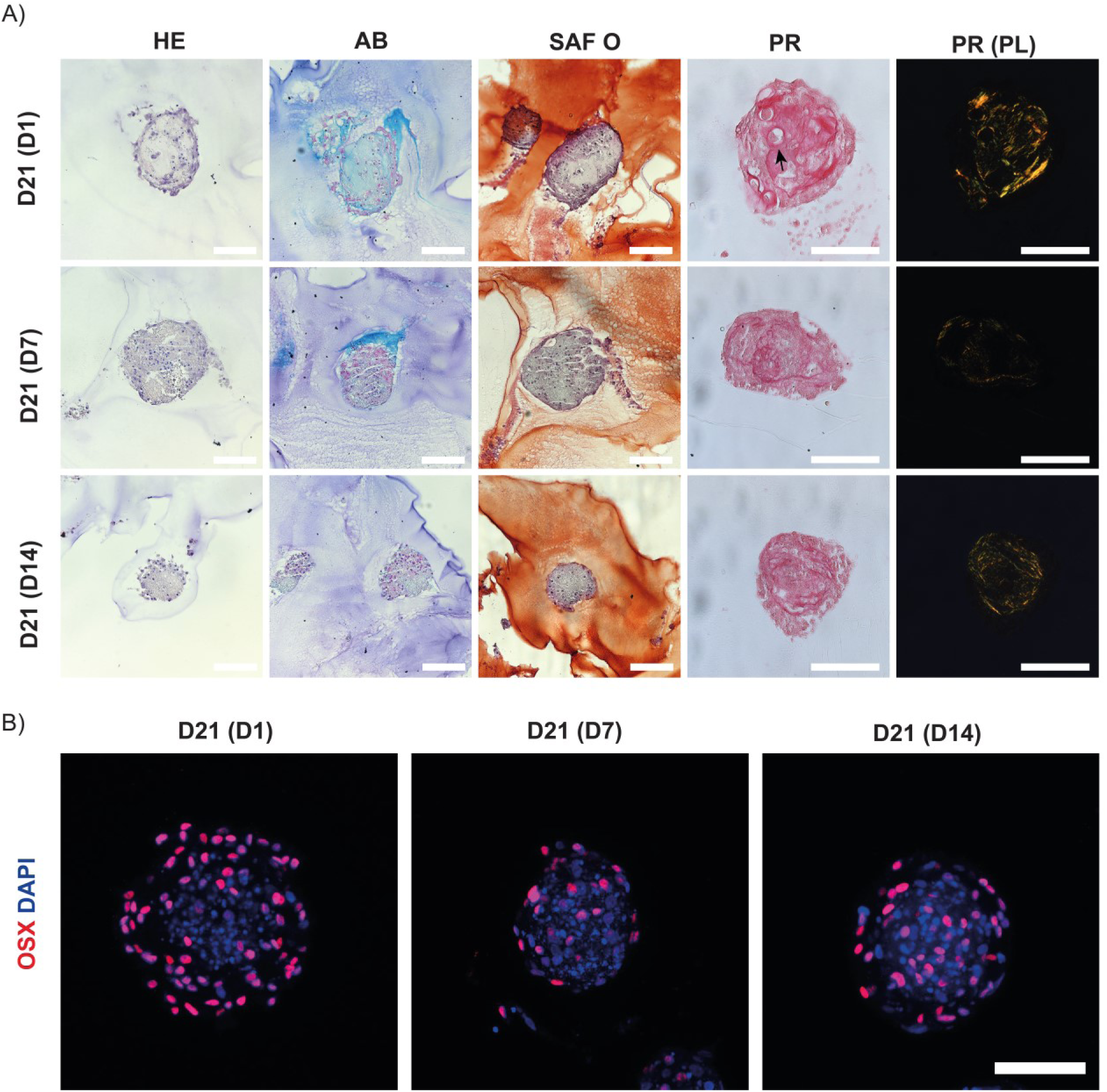
(A) Representative histological images and (B) representative maximum projection images of z-stacks obtained by confocal immunofluorescence of hPDC spheroids analyzed at day 21 after encapsulation on day 1 (D21 (D1)), day 7 (D21 (D7)), or day 14 (D21 (D14)). Histological stains include Hematoxylin and Eosin (HE), Alcian Blue (AB), Safranin O (SAF O), Picro-Sirius Red (PR), and PR under polarized light (PR(PL)). An example of a *lacuna* is highlighted with a black arrow. Immunofluorescence staining was performed for Osterix (OSX, red), with nuclei counterstained using DAPI (blue). Scale bars represent 100 µm.

On day 21, spheroids were immunostained for the transcription factor Osterix (OSX, Figure 2B). Positive staining for OSX was observed in all conditions, indicating the development of osteogenic differentiation and the existence of pre-hypertrophic chondrocytes. To confirm staining specificity, negative controls using a rabbit IgG isotype control and secondary antibody alone showed no detectable signal (Figure S1).

All encapsulation time points maintained high cell viability and supported comparable differentiation, as indicated by ECM deposition and Osterix expression. However, day 1 encapsulation resulted in more spheroid fusion and a tissue morphology that appeared more organized, with collagen fibers that were more distinctly visible under polarized light and the presence of *lacunae*. Based on these observations, day 1 was selected as the optimal encapsulation time point for downstream experiments.

### 3.2. Comparison of Differentiation Between Microwell-Cultured and HAMA-Encapsulated hPDC Spheroids in Hypertrophic Chondrogenic Medium: Cell Viability, Morphological Transitions and GAGs production

Next, we compared the degree of differentiation of hPDC spheroids cultured in microwell plates for 21 days in HCM versus spheroids encapsulated in HAMA after 1 day of microwell culture and maintained in HCM for an additional 20 days (Figure 3A).

**Figure 3.**
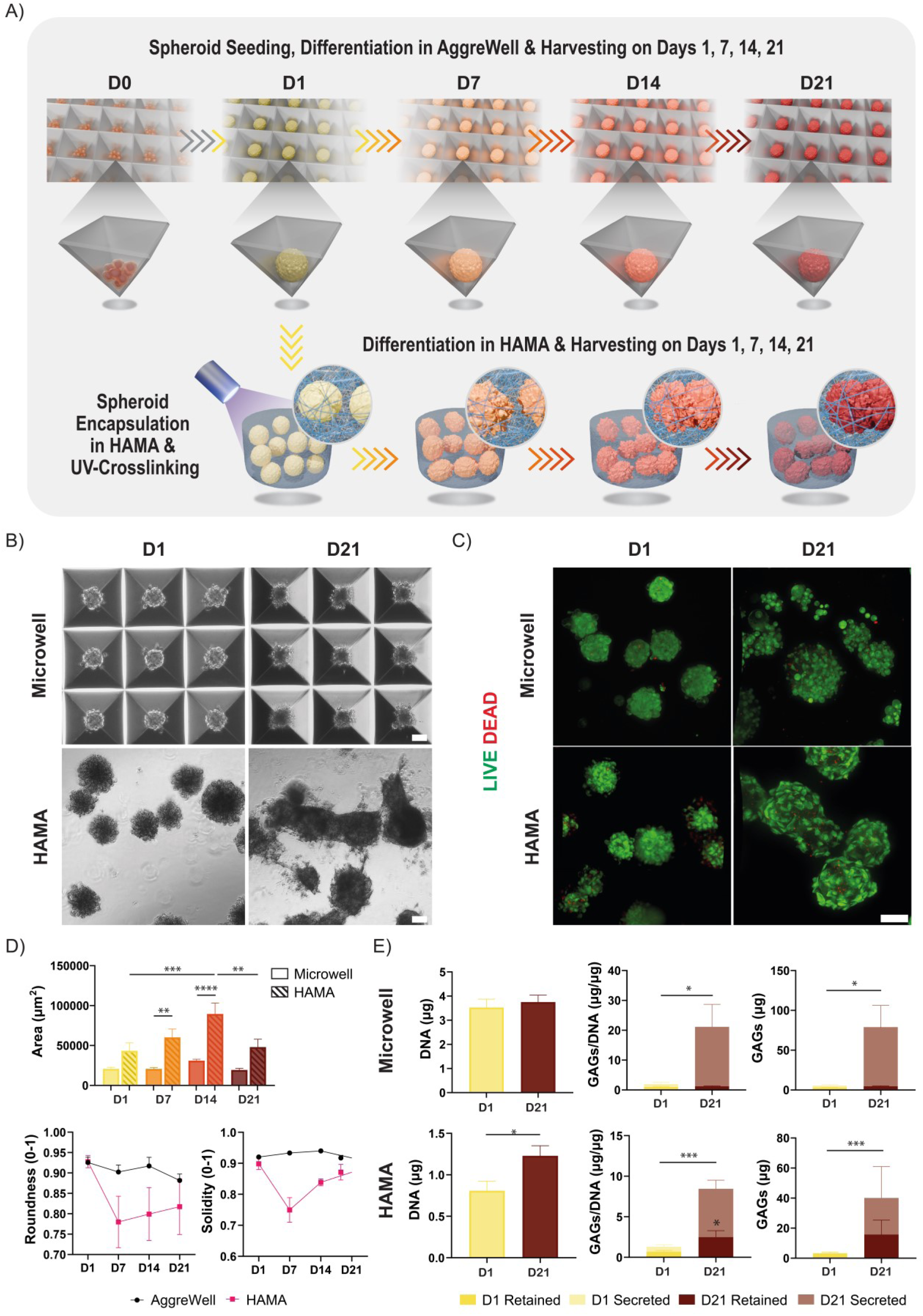
(A) Schematic representation of the experimental setup comparing hPDC spheroids cultured for 21 days in microwell plates versus those encapsulated in HAMA on day 1 and cultured for an additional 20 days in hypertrophic chondrogenic medium (HCM). (B) Representative bright-field images of spheroids differentiated in microwell plates or HAMA on day 1 and day 21. (C) Representative images of live-dead staining on day 1 and day 21. Live cells were stained with Calcein-AM (green), and dead cells were stained with EthD-1 (red). Scale bars represent 100 µm. (D) Quantification of spheroid cross-sectional area (top), roundness (bottom left), and solidity (bottom right) for both microwell and HAMA conditions at days 1, 7, 14, and 21. (E) Quantification of DNA content (left), GAGs/DNA (middle), and total GAGs content (right) for microwell (top) and HAMA (bottom) conditions on day 1 and day 21. For microwell condition a well was used (∼1200 spheroids) whereas for HAMA condition an encapsulated hydrogel (∼300 spheroids) was used. The GAG/DNA graph displays stacked bars representing both retained and secreted levels. Data are presented as mean ± SEM (*N* = 3). Statistical significance was determined using an unpaired t-test (\**p* < 0.05; \*\*\**p* < 0.001).

Bright-field images showed that spheroids encapsulated in HAMA were visibly larger and more irregular in shape compared to those cultured in microwell plates, with evidence of spheroid fusion by day 21 (Figure 3B). Live-dead staining performed on day 1 and day 21 demonstrated good cell viability in both conditions, indicating that neither the encapsulation nor the long-term culture affected cell survival (Figure 3C).

Quantification of spheroid cross-sectional area over time showed that HAMA-encapsulated spheroids were consistently larger than those cultured in microwell plates across all time points (Figure 3D, top). In the HAMA condition, spheroid area increased slightly from day 1 to day 7, and reached a peak at day 14, with significantly larger size compared to both day 1 and day 21. In contrast, spheroids in the microwell condition showed fluctuations in size over time, with an increase at day 14 followed by a decrease at day 21. However, these changes were not statistically significant. Comparison between conditions showed that HAMA-encapsulated spheroids were significantly larger than microwell-cultured spheroids at days 7 and 14. To assess changes in spheroid morphology, roundness and solidity were evaluated over time (Figure 3D, bottom). Roundness, which ranges from 0 to 1 and reflects how closely the shape approximates a perfect circle (1 being perfectly round), remained high in microwell-cultured spheroids throughout the culture period. In contrast, HAMA-encapsulated spheroids displayed a marked reduction in roundness at day 7, with partial recovery by day 21. Solidity, a measure of structural compactness calculated as the ratio of spheroid area to its convex hull area (with values closer to 1 indicating fewer surface irregularities), remained stable in spheroids cultured in microwells but decreased in HAMA-encapsulated spheroids at day 7 before increasing again by day 21.

Quantification of DNA content revealed no significant difference in DNA content between day 1 and day 21 for spheroids cultured in microwell plates (Figure 3E, top left). However, spheroids encapsulated in HAMA showed significantly higher DNA content at day 21, indicating enhanced proliferation or retention of cells in the 3D matrix (Figure 3E, bottom left). GAG content normalized to DNA showed a significant increase in secreted GAGs for both conditions by day 21, while retained GAGs showed significant difference in HAMA-encapsulated spheroids only (Figure 3E, middle). Total secreted GAG content was significantly higher for both conditions on day 21 compared to day 1 (Figure 3E, right).

To further analyze the morphological variability observed in microwell and HAMA conditions, unsupervised clustering was performed using principal component analysis (PCA) on image-derived features from phalloidin-stained confocal microscopy images. This analysis identified three distinct clusters (Figure 4A). Representative images of spheroids located along the PCA axes illustrate how variation along PC1 and PC2 captures distinct morphological trends (Figure 4B-C). Along PC2, higher values were associated with rounded actin distribution around cells within the spheroid, typical of early stages, while lower values corresponded to more elongated actin filaments indicative of cell fusion in the spheroid (Figure 4B). Along the PC1 axis, increasing values corresponded to more irregular, spiky morphologies, while lower values reflected compact, rounded spheroids (Figure 4C).

**Figure 4.**
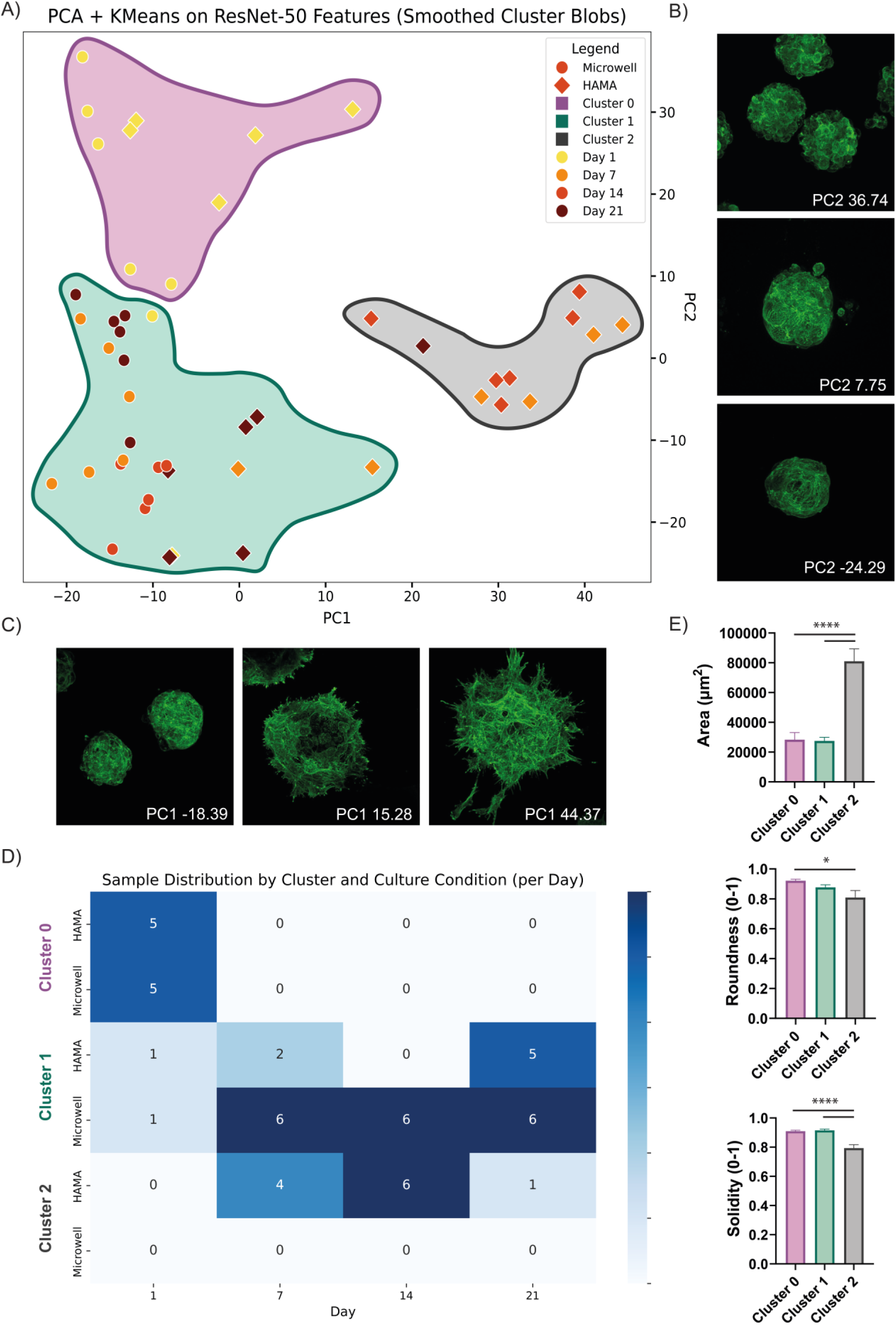
(A) Principal component analysis (PCA) plot of phalloidin-stained spheroids based on shape descriptors extracted from phalloidin-stained images. Colors indicate cluster identity; shapes and outlines denote culture condition and time point. (B) Representative phalloidin-stained images corresponding to low, intermediate, and high values along PC2. (C) Representative images corresponding to high, intermediate, and low values along PC1. (D) Heatmap of sample distribution by cluster, culture condition, and time point. (E) Quantification of cross-sectional area (top), roundness (middle), and solidity (bottom) per cluster. Data are shown as mean ± SEM (*N* = 3). Asterisks indicate significance from one-way ANOVA with Tukey’s post-hoc test (\**p* < 0.05, \*\*\*\**p* < 0.0001).

Sample distribution across clusters revealed clear trends in morphology linked to culture condition and culture duration (Figure 4D). Cluster 0 consisted entirely of day 1 spheroids, evenly split between microwell and HAMA. Cluster 1 included spheroids from both materials at later time points, particularly microwell-cultured spheroids from days 7, 14, and 21, and HAMA-encapsulated spheroids from day 21. In contrast, Cluster 2 was composed exclusively of HAMA-encapsulated spheroids, predominantly from days 7 and 14, indicating that the HAMA matrix promoted a unique morphological phenotype absent in microwell cultures.

Quantification of morphological parameters by cluster revealed that spheroids in Cluster 2 had significantly larger areas and lower solidity compared to Clusters 0 and 1, and significantly lower roundness relative to Cluster 0 (Figure 4E), supporting the morphological distinctions observed in the clustering analysis.

### 3.3. Comparison of Differentiation Between Microwell-Cultured and HAMA-Encapsulated hPDC Spheroids in Hypertrophic Chondrogenic Medium: Histology

To further investigate matrix composition and tissue organization, histological staining was performed on both microwell-cultured and HAMA-encapsulated spheroids harvested on days 1, 7, 14, and 21 post-spheroid formation. Figure 5 shows representative images of AB, SAF O, PR, and PR (PL) staining for microwell and HAMA samples. HE staining is provided in Figure S2.

**Figure 5.**
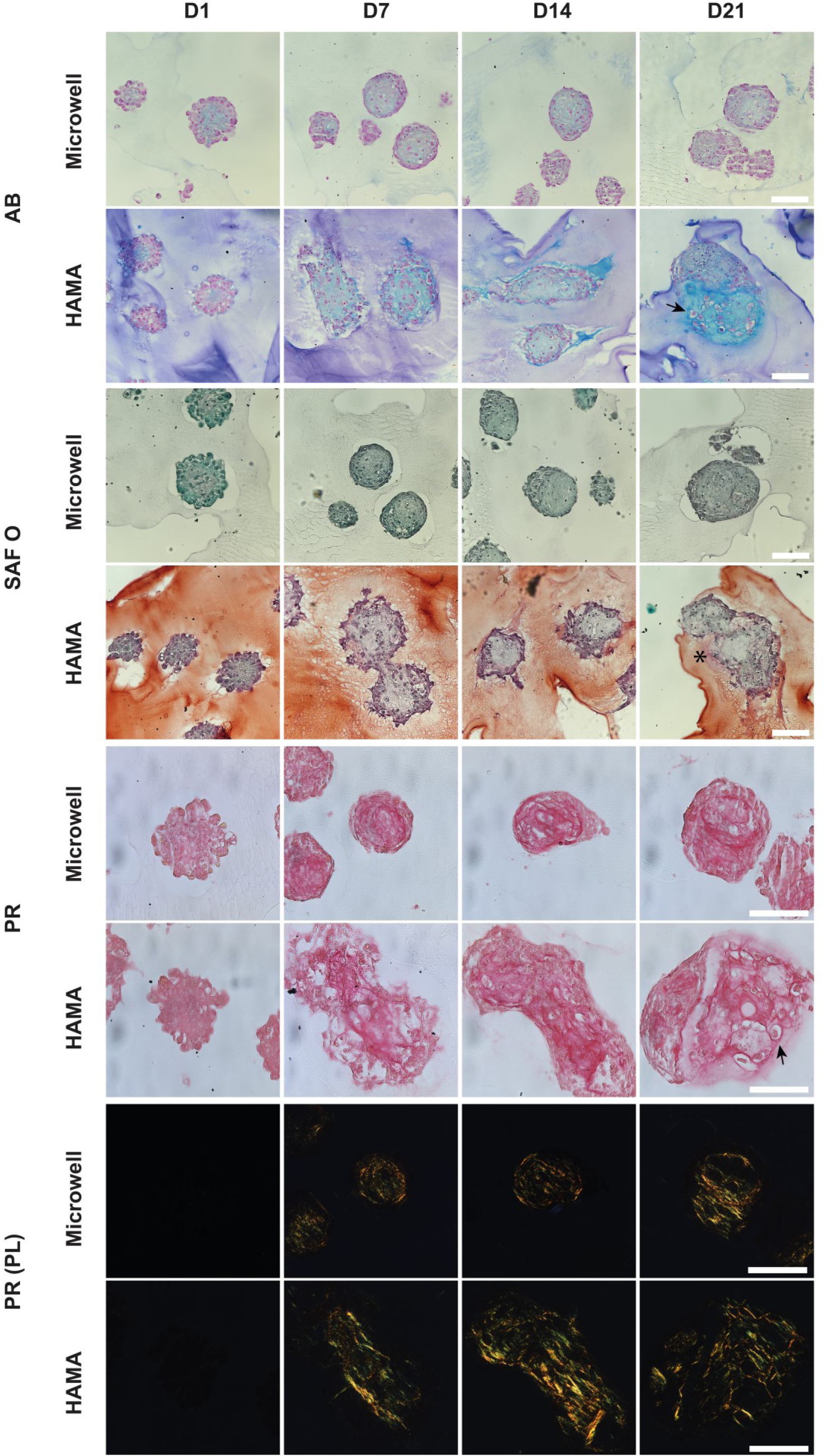
Representative images of histological staining of microwell-cultured and HAMA-encapsulated hPDC spheroids harvested on days 1, 7, 14, and 21. Stainings include Alcian Blue (AB), Safranin O (SAF O), Picro-Sirius Red (PR) and PR under polarized light (PR(PL)). Examples of *lacunae* are indicated with black arrows. SAF O positivity is highlighted with an asterisk. Scale bars are 100 µm.

In both conditions, AB positivity, indicative of GAG presence, was visible from day 1. From day 14, AB staining was also around the HAMA-encapsulated spheroids, which may reflect GAG retention within the hydrogel. By day 21, the HAMA samples also showed the presence of *lacunae*, which were absent in the microwell samples (Figure 5). SAF O positivity, which highlights sulfated proteoglycans, was detected in HAMA-encapsulated spheroids on day 21 (Figure 5, Figure S3). PR staining showed collagen presence in both conditions. These fibers began to be visible under polarized light from day 7 onwards, with more pronounced collagen signal and apparent alignment in the HAMA samples.

### 3.4. Progression of Osteogenic and Chondrogenic Marker Expression in Microwell-Cultured and HAMA-Encapsulated Spheroids

To evaluate the expression over time of key chondrogenic and osteogenic markers, immunostaining for SRY-box transcription factor 9 (SOX9), Osterix (OSX), and Runt-related transcription factor 2 (RUNX2) was performed on days 1, 7, 14, and 21. Figure 6 presents representative immunostaining images for both microwell-cultured and HAMA-encapsulated spheroids.

**Figure 6.**
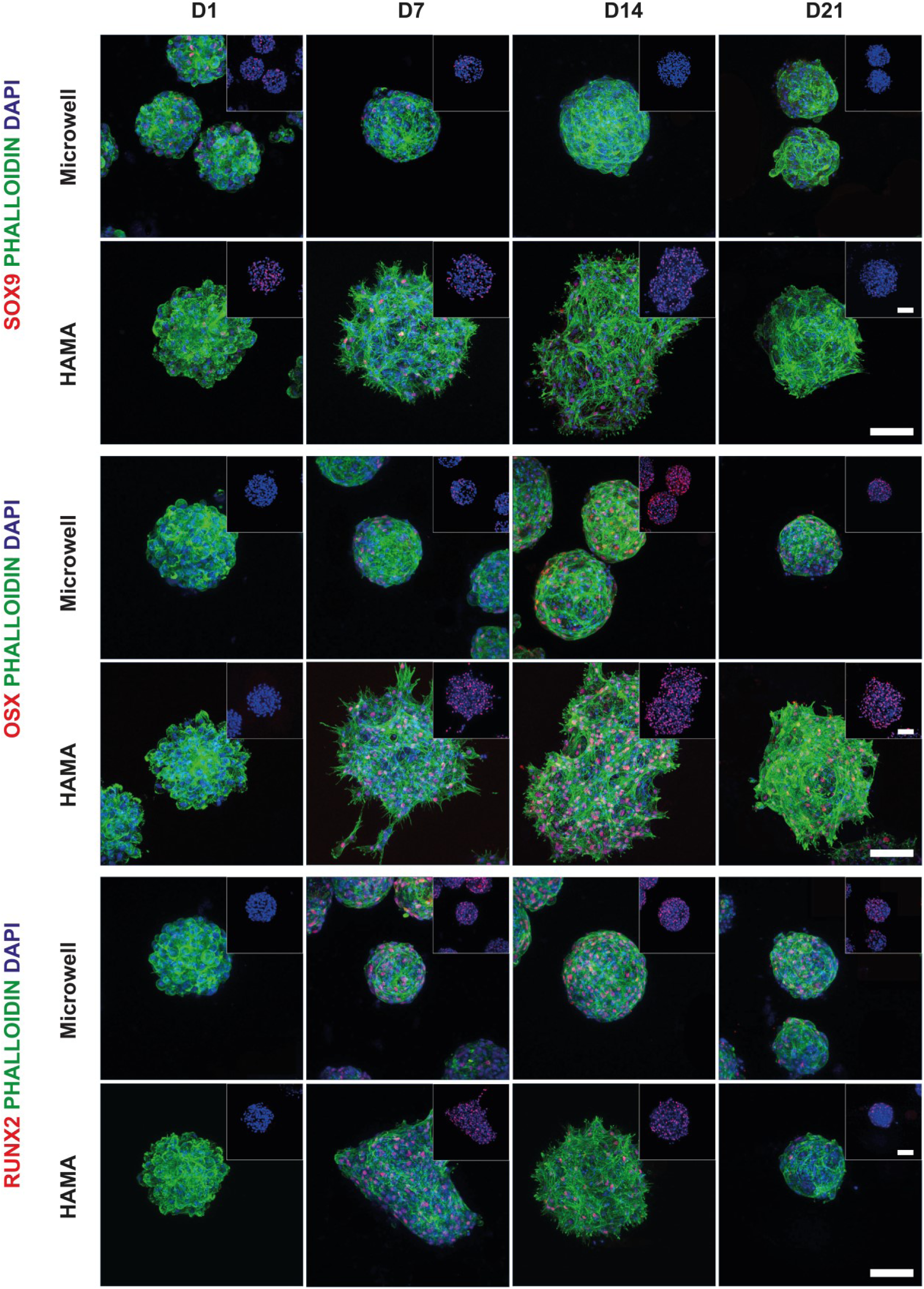
Representative maximum projection images of z-stacks obtained by confocal immunofluorescence staining for SRY-box transcription factor 9 (SOX9, red), Osterix (OSX, red), and Runt-related transcription factor 2 (RUNX2, red) in microwell-cultured and HAMA-encapsulated spheroids at days 1, 7, 14, and 21. Cell nuclei were stained with DAPI (blue) and actin filaments were stained with phalloidin (PHALLOIDIN, green). Insets in the top right corner show the sample stained in blue (DAPI) and red (OSX, SOX9, or RUNX2). Scale bars represent 100 µm.

SOX9 expression, a marker of early chondrogenesis, followed a different pattern in the two conditions. In microwell-cultured spheroids, SOX9 was positive only at days 1 and 7, whereas in HAMA-encapsulated spheroids, it remained positive through days 1, 7, and 14. OSX, associated with hypertrophic chondrocytes and osteogenic commitment, was detected starting from day 7 in both conditions. RUNX2, a critical regulator of chondrocyte hypertrophy and osteogenic differentiation, was expressed on days 7, 14, and 21 in both conditions. To confirm staining specificity, negative controls using a rabbit IgG isotype control and secondary antibody alone showed no detectable signal (Figure S4). These results suggest that cells in both culture systems gradually shift from an early chondrogenic state toward a more hypertrophic phenotype. The changes in SOX9, OSX, and RUNX2 over time reflect a pattern similar to early steps seen in endochondral ossification.

### 3.5. Gene Expression Analysis of Key Chondrogenic and Osteogenic Markers in Microwell-Cultured and HAMA-Encapsulated Spheroids

RT-qPCR analysis was performed to assess the expression of key chondrogenic and osteogenic genes over time in both microwell-cultured and HAMA-encapsulated spheroids. The primers used included *ACAN* (Aggrecan), *COL2A1* (type II collagen), *COL1A1* (type I collagen), *COL10A1* (type X collagen), *OPN* (osteopontin), *ALPL* (alkaline phosphatase), and *IBSP* (Integrin Binding Sialoprotein).

In microwell-cultured spheroids (Figure 7A), *ACAN* expression peaked at day 14, indicating a mid-point increase in proteoglycan synthesis. *COL2A1*, a marker of cartilage matrix production, showed its highest fold change on day 21. *COL1A1* expression was highest at days 7 and 14, while *COL10A1*, a marker of hypertrophic chondrocytes, and *OPN*, associated with mineralization, both increased over time and peaked at day 21. *ALPL*, involved in early osteogenic differentiation, showed a general increasing trend, although no significant differences were observed across time points. *IBSP*, another osteogenic marker, showed an increase over time, peaking at day 21.

**Figure 7.**
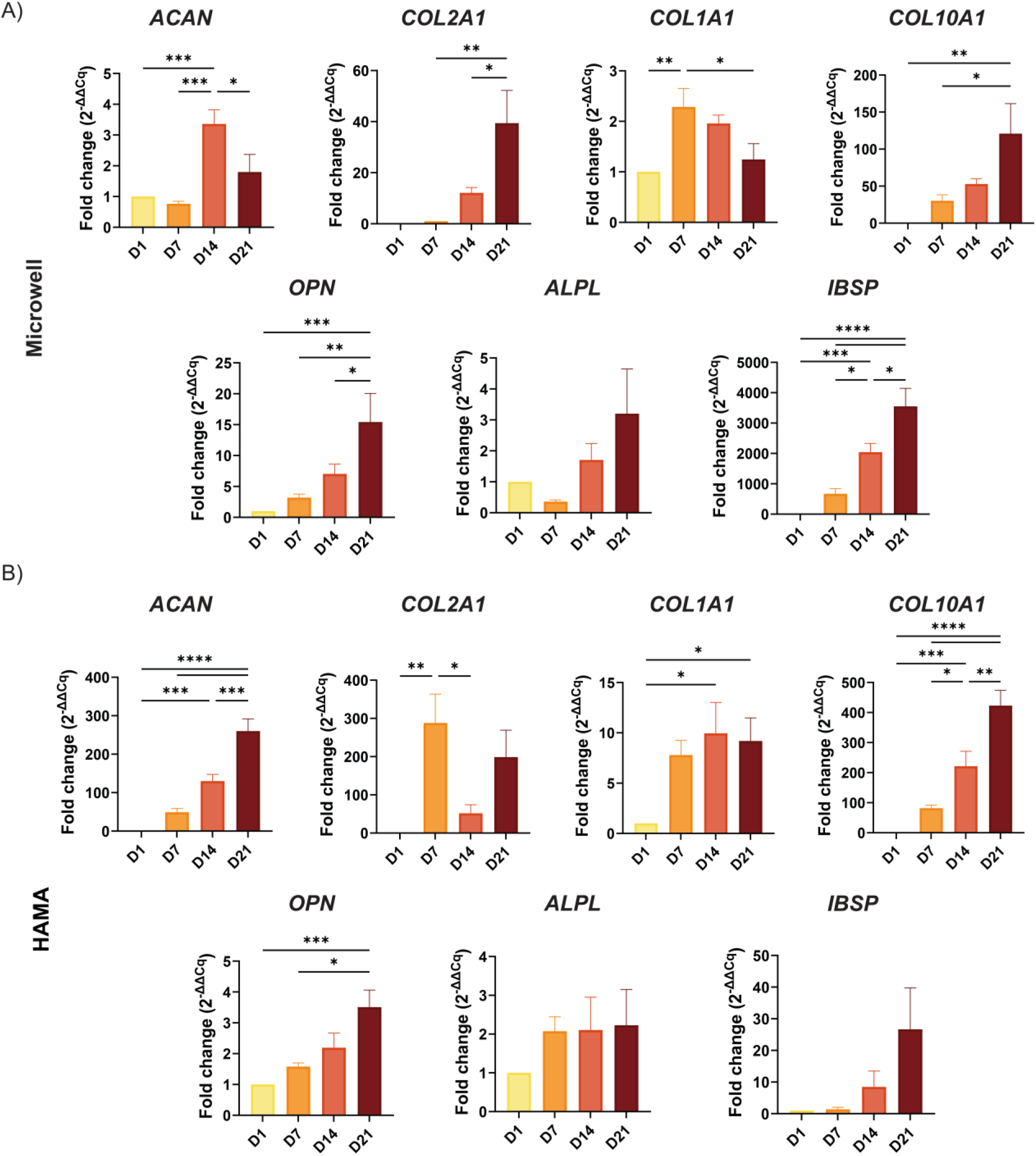
Quantification of mRNA transcript of key chondrogenic and osteogenic markers in microwell-cultured (A) and HAMA-encapsulated spheroids (B) at different time points, normalized to day 1. For *COL2A1* in the microwell condition, day 7 was used as the reference because transcripts were not detected on day 1. Data are presented as mean ± SEM (*N* = 3). Statistical significance was determined using one-way ANOVA followed by Tukey’s post-hoc multiple comparison test (\**p* < 0.05; \*\**p* < 0.01); \*\*\**p* < 0.001; \*\*\*\**p* < 0.0001).

In HAMA-encapsulated spheroids (Figure 7B), *ACAN* expression steadily increased until day 21. *COL2A1* expression peaked on day 7, with no significant difference on day 21, indicating early cartilage matrix production. *COL1A1* expression was significantly higher on days 14 and 21 compared to day 1, reflecting the later stages of matrix remodeling. *COL10A1* expression increased progressively over time, peaking at day 21, indicating the presence of hypertrophic chondrocytes in the later stages of culture. *OPN* expression increased and peaked on day 21, reflecting enhanced mineralization. *ALPL* expression showed no significant differences over time. *IBSP* expression increased over time but did not show significant differences across the time points.

### 3.6. hPDC Spheroid High Density Bioprinting and Differentiation in Hypertrophic Chondrogenic Medium

The comparative analysis of encapsulated and microwell-cultured spheroids showed that HAMA provided a supportive three-dimensional environment that enhanced spheroid fusion, promoted extracellular matrix deposition, and supported the formation of a more mature cartilage-like tissue, including the appearance of *lacunae*. Gene expression and immunostaining confirmed progression toward hypertrophic phenotype in both conditions, with differences in the timing of some key markers. Based on these findings, we next explored whether the benefits observed with HAMA encapsulation could be maintained in bioprinted constructs, which offer improved structural organization and potential for clinical scale-up.

Initially, spheroids were bioprinted using a 250 µm nozzle, which resulted in high cell mortality and lack of differentiation, as evidenced by live-dead staining and the absence of OSX positivity on day 21 (Figure S5). Given these challenges, bioprinting was repeated with a 400 µm nozzle. However, the low viscosity of the bioink led to limited extrusion control and unstable printing. To address this, a partial crosslinking strategy was introduced before printing. To determine optimal crosslinking conditions, photo-rheology experiments were performed (Figure S6A), in which a solution containing 2% w/v HAMA and 0.01% w/v LAP in PBS was exposed to UV light for 30, 40, 50, or 60 seconds. As expected, there was an increase in G’ with increasing crosslinking time (Figure S6B). Based on these findings, a 40-second pre-crosslinking time was selected, as this provided a gel-like consistency suitable for bioprinting. A bioprinting trial comparing 30 and 40 second pre-crosslinking times showed higher shape fidelity and printability with 40 seconds (Figure S6C).

After optimizing the bioprinting conditions, spheroids were bioprinted on day 1 using a HAMA-based bioink. The spheroid-laden bioink was loaded into a syringe (Figure S7A) and extruded using a pattern of connected circles (Figure S7B, left), visible in the one-layer construct on day 1 (Figure S7B, middle). By day 21, in the four-layer construct, an increased opacity and spheroid fusion was observed which suggested an increase in matrix deposition and differentiation (Figure S7B, right). In some specific areas, where spheroids were aligned, partial or complete fusion was observed on day 21 (Figure S7C).

The bioprinting process and subsequent culture setup are shown schematically in Figure 8A. Bright-field images taken on days 1 and 21 showed clear evidence of spheroid fusion, resulting in larger spheroids over time (Figure 8B). Live-dead staining confirmed good cell viability immediately after bioprinting (day 1) and on day 21, indicating that both the bioprinting process and long-term culture in HCM supported cell survival (Figure 8C).

**Figure 8.**
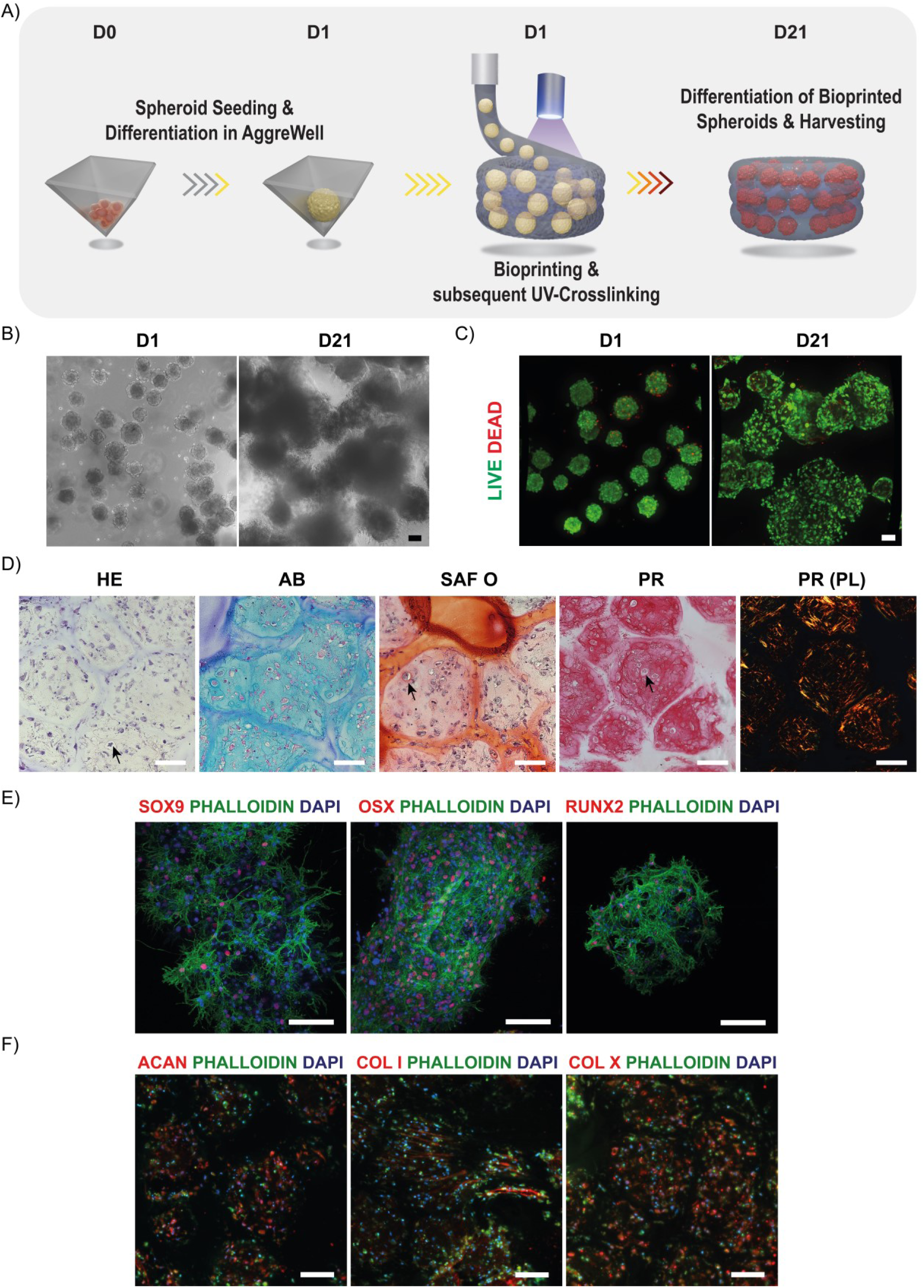
(A) Schematic of the bioprinting process: hPDC spheroids were bioprinted in HAMA on day 1 and cultured in hypertrophic chondrogenic medium until day 21. (B) Bright-field images on days 1 and 21. (C) Live-dead staining at days 1 and 21. Live cells were stained with Calcein-AM (green), and dead cells were stained with EthD-1 (red). (D) Representative images of histological staining on day 21 using Hematoxylin and Eosin (HE), Alcian Blue (AB), Safranin O (SAF O), Picro-Sirius Red (PR) and PR under polarized light (PR(PL)). Examples of *lacunae* are highlighted with black arrows. (E) Representative maximum projection images of z-stacks obtained by confocal immunofluorescence staining for SRY-box transcription factor 9 (SOX9, red), Osterix (OSX, red), and Runt-related transcription factor 2 (RUNX2, red) on day 21. Cell nuclei were stained with DAPI (blue) and actin filaments were stained with phalloidin (PHALL, green). (F) Representative immunostaining images of sectioned samples stained for Aggrecan (ACAN, red), collagen type I (COL I, red), and collagen type X (COL X, red) at day 21. Cell nuclei were stained with DAPI (blue) and actin filaments were stained with phalloidin (PHALLOIDIN, green). Scale bars represent 100 µm.

Histological staining on day 21 demonstrated strong positivity for AB, SAF O, and PR (Figure 8D). AB staining revealed high GAG deposition, indicative of cartilage matrix formation. SAF O staining highlighted the presence of GAGs within the fused spheroids, seen as a red-orange color. PR staining showed strong collagen fiber formation, which was further confirmed under polarized light, where birefringence indicated thick, well-organized collagen fibers. Additionally, *lacunae*, indicative of cartilage matrix maturation, were visible in the stainings, with examples indicated by arrows.

Immunofluorescence staining on day 1 showed SOX9 positivity, with no detectable expression of OSX or RUNX2, consistent with a chondrogenic phenotype, similar to that observed in encapsulated spheroids (Figure S7D). Immunostaining results on day 21 (Figure 8E) revealed positive staining for OSX, SOX9, and RUNX2, markers of osteogenic and chondrogenic differentiation, confirming the progression toward these lineages. OSX and RUNX2 positivity indicated osteogenic differentiation and hypertrophy, respectively, while SOX9 highlighted chondrogenesis. Additional immunostaining showed positive expression of aggrecan (ACAN), collagen type X (COL X), and collagen type I (COL I) supporting cartilage matrix production, hypertrophy, and osteogenic differentiation, respectively (Figure 8F). The evidence from histology and immunostaining confirmed the development of hypertrophic cartilage within the bioprinted constructs. To confirm staining specificity, negative controls using a rabbit IgG isotype control and secondary antibody alone showed no detectable signal (Figure S8).

Overall, the results demonstrated that encapsulating hPDC spheroids in HAMA hydrogels at an early differentiation timepoint enhanced cell viability, promoted fusion, and supported cartilage-like tissue maturation, as indicated by increased matrix deposition and *lacunae* formation. When compared to culture in microwell plates, encapsulated spheroids showed distinct morphological changes, higher DNA content, and matrix characteristics consistent with more advanced stages of chondrogenic differentiation. Across conditions, gene expression and immunostaining confirmed progression toward hypertrophic and osteogenic phenotype. Finally, extrusion bioprinting of HAMA-encapsulated spheroids preserved viability and differentiation capacity in spatially patterned constructs, supporting the potential of this approach for scalable tissue engineering applications.

## 4. Discussion

Scaffold-free spheroid systems are highly valuable in developmental BTE because they closely mimic the early biological processes of bone formation, such as cell-cell interactions, self-assembly, and condensation [31]. However, their clinical utility is constrained by inherent limitations in scalability and construct integrity [32]. Incorporating biomaterials can overcome these issues by offering a stabilizing framework that supports spatial organization and mechanical cohesion, reduces the amount of cells required but also delivers instructive microenvironmental cues that promote tissue-specific differentiation [33]. Our goal was to bioprint hPDC spheroids at high density within a HA matrix to generate clinically relevant and scalable hypertrophic cartilage implants for BTE applications.

3D bioprinting shows significant potential for developing complex, functional tissue constructs for clinical applications [34]. A key advantage of extrusion-based bioprinting lies in its ability to deposit spheroid-based bioinks through a nozzle, forming continuous, densely packed cellular structures. This bioprinting method is useful for high-cell-density bioprinting, as it allows processing several living building blocks into relatively large living structures. It also supports scalability and efficiency by allowing individual spheroids to fuse directly on the receiving substrate [24, 35]. Additionally, the ability of hydrogels to support large tissue constructs with fewer spheroids presents a major scalability advantage. Hydrogels fill interstitial spaces and facilitate cellular infiltration and ECM deposition, allowing for the formation of sizable grafts with a lower total spheroid requirement. This significantly reduces cell sourcing demands and spheroid fabrication costs, which are often bottlenecks in large-scale tissue engineering.

Among candidate biomaterials for the bioink, HA, a major component of the cartilage ECM, was of special interest due to its role in activating endochondral ossification pathways, such as TGF-β, BMP, and Wnt in MSCs and chondrocytes, and facilitating cellular condensation and ECM deposition [25–27]. Furthermore, a HAMA-based bioink has significant translational advantages for treating long bone defects because HA-based products are already widely used in approved clinical applications with established safety, biocompatibility, and regulatory precedents [36, 37]. Its familiarity to regulatory agencies and availability in GMP-grade simplifies approval and clinical translation, especially in comparison to more novel or less-tested materials.

Achieving spheroid fusion is particularly relevant because it mimics the mesenchymal condensation phase of endochondral ossification, a critical early step in long bone development [38, 39]. By recapitulating this process *in vitro*, fused spheroids can initiate the chondrogenic and hypertrophic cascade, ultimately leading to the formation of a cartilage template that supports vascular invasion and ossification upon implantation. We also investigated the impact of encapsulation timing on spheroid fusion and differentiation. Previous studies with hPDC spheroids in material-free conditions showed less fusion capacity with increased maturation stage [38, 40]. Consistent with this, our results showed that early encapsulation of hPDC spheroids after one day of cell aggregation significantly improved fusion capacity and increased DNA content by day 21, likely due to greater mobility and minimal ECM, both of which are important for fusion. To further promote fusion, a low concentration HAMA hydrogel formulation (2% w/v) with a low stiffness (∼1 kPa) and a low methacrylation degree was used [29, 41, 42]. Additionally, high spheroid density reduced the inter-spheroid distance promoting efficient contact-mediated fusion. The higher DNA content observed may indicate enhanced cell proliferation or improved cell retention within the HAMA hydrogel, potentially due to a supportive environment that helps cell-cell and cell-matrix interactions. Having a structure around spheroids has previously been reported to retain non-spheroid forming cells that are normally expulsed into the culture media in microwell-cultured spheroids [43].

Even though early encapsulation promoted spheroid growth and fusion, the GAGs synthesis remained consistent, indicating that the differentiation capacity is maintained regardless of when the spheroids were encapsulated. High cell viability across all conditions shows the cytocompatibility of the HAMA hydrogel and the encapsulation process. Histological staining revealed that spheroids encapsulated on day 1 exhibited the presence of *lacunae* by day 21, an indicator of chondrocyte maturation and cartilage matrix development, showing progression toward hypertrophic cartilage formation [44, 45]. The absence of SAF O staining across all encapsulation time points, despite positive AB staining, suggests the presence of non-sulfated GAGs or early-stage proteoglycans, reflecting an immature cartilage matrix. However, increased collagen deposition and the presence of OSX, expressed by pre-hypertrophic chondrocytes and osteoblasts, indicate progression toward osteogenic differentiation, highlighting its role in late chondrogenesis and the shift to osteogenesis [46–48].

To contextualize the effects of encapsulation and matrix environment on spheroid behavior, we compared HAMA-encapsulated spheroids to those cultured in commercially available microwell plates. These comparative analyses between hPDC spheroids cultured in microwell plates and those encapsulated in HAMA showed the advantages of the 3D hydrogel environment. The increased size of HAMA-encapsulated spheroids reflects not only cellular proliferation but also dynamic fusion and cell outgrowth, likely facilitated by permissive matrix interactions. Similar behaviors have been observed in soft gelatin-based hydrogels, where embedded spheroids sprout and interconnect [49, 50], especially when early-stage spheroids are used [51].

Morphological analyses supported this dynamic behavior. HAMA-encapsulated spheroids underwent reduced roundness and solidity, reflecting a remodeling phase driven by fusion dynamics and cell-matrix interactions. Spiky, protrusive morphologies were present, consistent with cell migration and cytoskeletal reorganization at the spheroid periphery. Similar protrusions, and inter-spheroid bridging structures have been reported in Matrigel-embedded stem cell spheroids cultured under controlled spatial arrangements [52].

The presence of *lacunae* in HAMA-encapsulated hPDC spheroids, compared to their absence in material-free conditions, suggests that the surrounding matrix may play a role in promoting structural changes associated with chondrocyte hypertrophy. While the precise mechanisms remain unclear, one possible explanation could be the contribution of hyaluronan in the HAMA hydrogel. Pavasant *et al.* demonstrated that hyaluronan facilitates lacunar enlargement in the growth plate by attracting water and exerting swelling pressure on the pericellular matrix [53]. A similar mechanism may occur in our encapsulated spheroids, where the HAMA hydrogel provides a hyaluronan-rich environment that could influence cellular behavior and matrix remodeling. However, further studies would be necessary to confirm this hypothesis and understand HAMA’s specific effects on hPDC spheroid differentiation. GAGs quantification also revealed that the HAMA matrix helped retain some of the secreted GAGs within the hydrogel and around the spheroids.

The overall expression profiles of chondrogenic and osteogenic markers indicate that both microwell-cultured and HAMA-encapsulated spheroids are capable of progressing through some of the key stages of endochondral differentiation, but on slightly different timelines [54]. While spheroids in both conditions first went into chondrogenic differentiation, the immunostaining results showed prolonged SOX9 positivity in HAMA-encapsulated spheroids. This suggests that the hydrogel provides a microenvironment conducive to maintaining a chondrogenic phenotype for a more extended period [55]. SOX9 activates collagen type X transcription in hypertrophic chondrocytes and inhibits premature osteogenic commitment [56]. The steady increase in *ACAN* expression, indicative of ongoing proteoglycan deposition, further supports this. Maintaining an extended chondrogenic phase in some of the cells within the spheroids, with prolonged *SOX9* expression and increased GAG production and deposition, could lead to faster and more homogeneous new bone formation *in vivo* after implantation by providing a more robust cartilage template for remodeling [57–60].

RUNX2 is a key transcription factor for bone formation and osteoblast differentiation. Osterix is an essential transcription factor for endochondral ossification that works downstream of RUNX2 [61, 62]. Microwell-cultured and HAMA-encapsulated spheroids were positive for both markers from day 7, indicating differentiation towards a hypertrophic and osteoblastic phenotype. Similar trends in the expression of *COL10A1* (hypertrophy marker), *OPN* (pre- and osteoblast marker), and *IBSP* (osteoblast marker) confirm that the hypertrophy and shift towards osteoblastic differentiation also occur in the encapsulated spheroids [63–66].

Bioprinting hPDC spheroids at high density resulted in robust cartilage-like tissue formation, as demonstrated by high cell viability, significant GAG and GAGs deposition, and organized collagen fibers. The presence of *lacunae* indicated advanced chondrogenesis, and immunostaining for SOX9, RUNX2, and OSX confirmed progression toward a hypertrophic and osteogenic phenotype. The nozzle size had to be increased to avoid cell death caused by shear stress during extrusion, this often leads to a more random deposition of spheroids [67]. However, partially crosslinking the ink before adding the spheroids helped with aligning them in some regions. Although precise spatial control was limited, sporadic “tube-like” structures were formed by day 21 when spheroids were aligned closely (Figure S7C). This was not observed in the bulk encapsulation of hPDC spheroids, so we hypothesized that the extrusion direction of the partially crosslinked bioink contributed to spheroid orientation or helped with the spheroids being extruded one by one.

While our findings are promising, a few limitations should be acknowledged. First, the *in vitro* nature of our study does not fully capture the complexities of the *in vivo* environment, where factors such as vascularization, mechanical loading, and immune responses significantly influence tissue development and integration. Second, the lack of precise control over the exact positioning of spheroids within the bioprinted constructs which may affect the consistency and functionality of the engineered tissue, regardless of their fusion or differentiation state. Therefore, using emerging bioprinting approaches, such as embedded bioprinting, laser-assisted bioprinting, and aspiration-assisted bioprinting, offer potential solutions by improving spheroid positioning and spatial resolution [68–71]. These methods have demonstrated improved structural integrity and deposition precision but require further optimization to enhance printing efficiency and scalability, as well as reduce the risk of spheroid rupture [24].

## 5. Conclusions

This study demonstrates the feasibility of bioprinting bioinks composed of high-density hPDC spheroids and HAMA as a scalable strategy for engineering hypertrophic cartilage constructs for bone regeneration. The observed spheroid fusion, ECM deposition, and progressive chondrogenic-to-osteogenic differentiation emphasize the potential of this approach for large bone defect repair. Notably, this work highlights that HAMA can actively support spheroid fusion and hypertrophic cartilage formation; key factors in translating bioprinted constructs into clinically relevant bone grafts. Future research should focus on integrating vascularization strategies, refining deposition techniques for better spatial resolution, and validating the functional performance of these constructs *in vivo*.

## Acknowledgments

The authors acknowledge the Tissue Engineering Lab, Skeletal Biology and Engineering Research Centre, Department of Development and Regeneration, KU Leuven, and Prometheus, the Leuven R&D division of Skeletal Tissue Engineering of KU Leuven, for providing hPDCs and sharing valuable expertise. We thank Marina Maréchal for her guidance and support regarding hPDC differentiation, and Inge Van Hoven and Samuel Ribeiro for training and technical assistance in hPDC culture, spheroid generation, and differentiation.

## Data availability statement

The data that support the findings of this study are available from the corresponding author upon reasonable request.

## Ethics approval statement

All procedures involving human periosteal tissue were approved by the Ethical Committee for Human Medical Research (KU Leuven), and informed consent was obtained from all patients (approval number: ML7861).

