## Supplementary material for "Packed for Ossification: High-Density Bioprinting of hPDC Spheroids in HAMA for Endochondral Ossification"

**Table S1.** qPCR primer sequences. List of primers used and their sequences (5' to 3').

| Primer name | Primer sequence (5' → 3') |
|---|---|
| PSMB2_Fw | GCTGCCAGGTAGTCCATGTAA |
| PSMB2_Rv | CGAAACCTGGCTGACTGTCT |
| REEP5_Fw | AGGTCAGCCACTGGGTATCA |
| REEP5_Rv | CCTCTCTCCTCTGCAACCTG |
| COL2A1_Fw | GGCTTCCATTTCAGCTATGG |
| COL2A1_Rv | AGCTGCTTCGTCCAGATAGC |
| OPN_Fw | TGAAACGAGTCAGCTGGATG |
| OPN_Rv | TGAAATTCATGGCTGTGGAA |
| COL10A1_Fw | ACGATACCAAATGCCCACAG |
| COL10A1_Rv | GTGGACCAGGAGTACCTTGC |
| ALPL_Fw | GCTTCAAACCGAGATACAAGCA |
| ALPL_Rv | GCTCGAAGAGACCCAATAGGTAGT |
| ACAN_Fw | GTCTCACTGCCCAACTAC |
| ACAN_Rv | GGAACACGATGCCTTTCAC |

| | |
|---|---|
| IBSP_Fw | GGCAGTAGTGACTCATCCGAAG |
| IBSP_Rv | GAAAGTGTGGTATTCTCAGCCTC |
| COL1A1_Fw | GAGGGCCAAGACGAAGACATC |
| COL1A1_Rv | CAGATCACGTCATCGCACAAC |

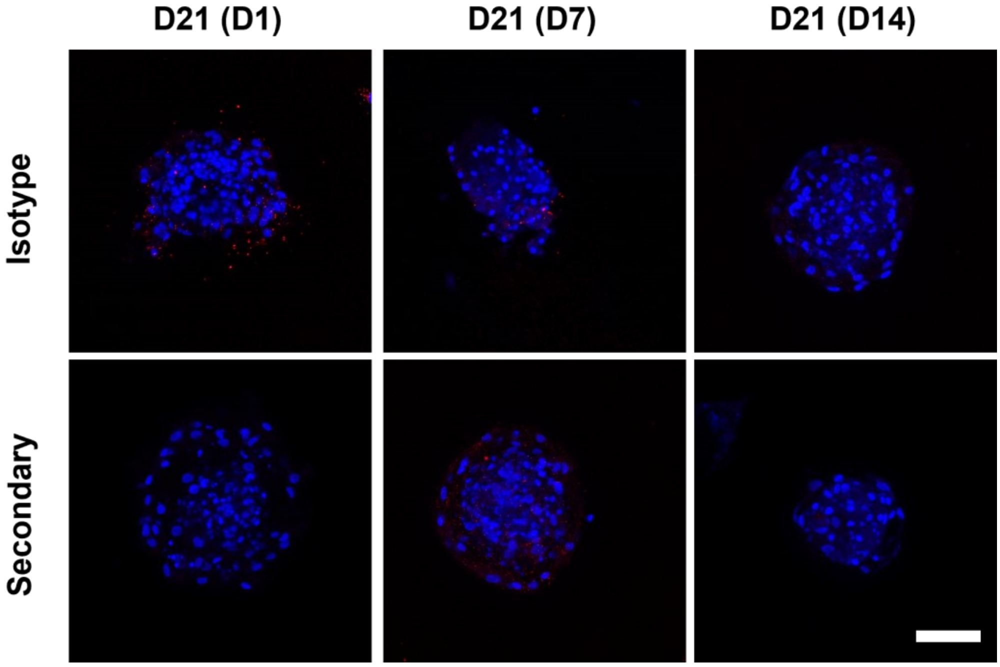


**Figure S1.** Representative maximum projection images of z-stacks obtained by confocal immunofluorescence staining of negative controls for hPDC spheroids on day 21 and encapsulated on day 1, 7 or 14. Negative controls included samples where an IgG isotype control antibody was used (top row) or where no primary antibody was added (bottom row). Cell nuclei were stained with DAPI (blue). Scale bar is 100 µm.

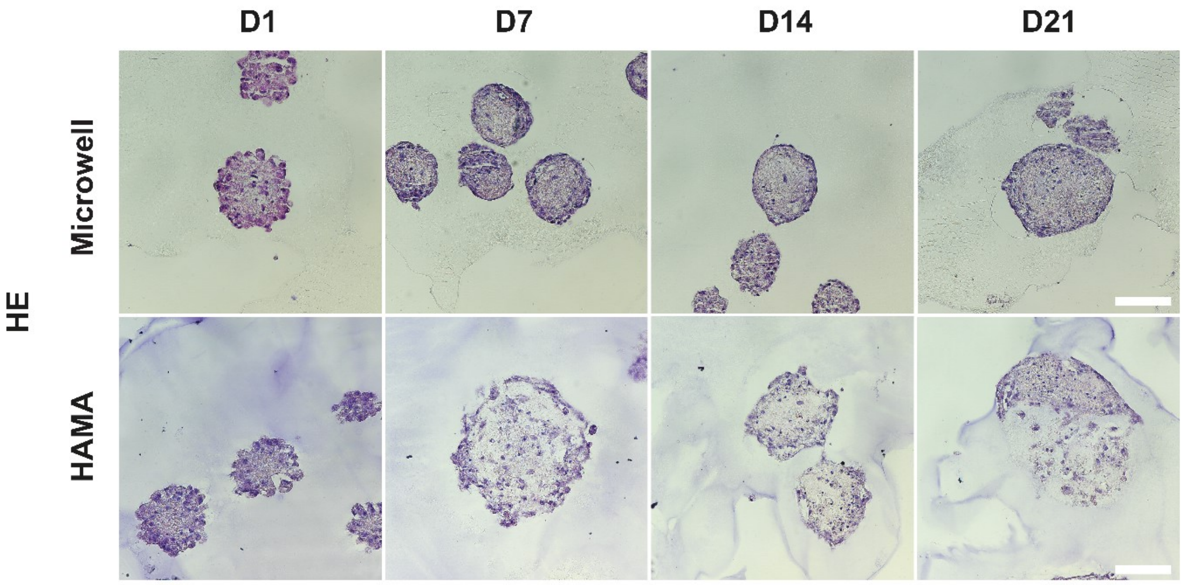


**Figure S2.** Representative images of HE staining of microwell-cultured and HAMA-encapsulated hPDC spheroids harvested on days 1, 7, 14, and 21. Scale bars are 100 µm.

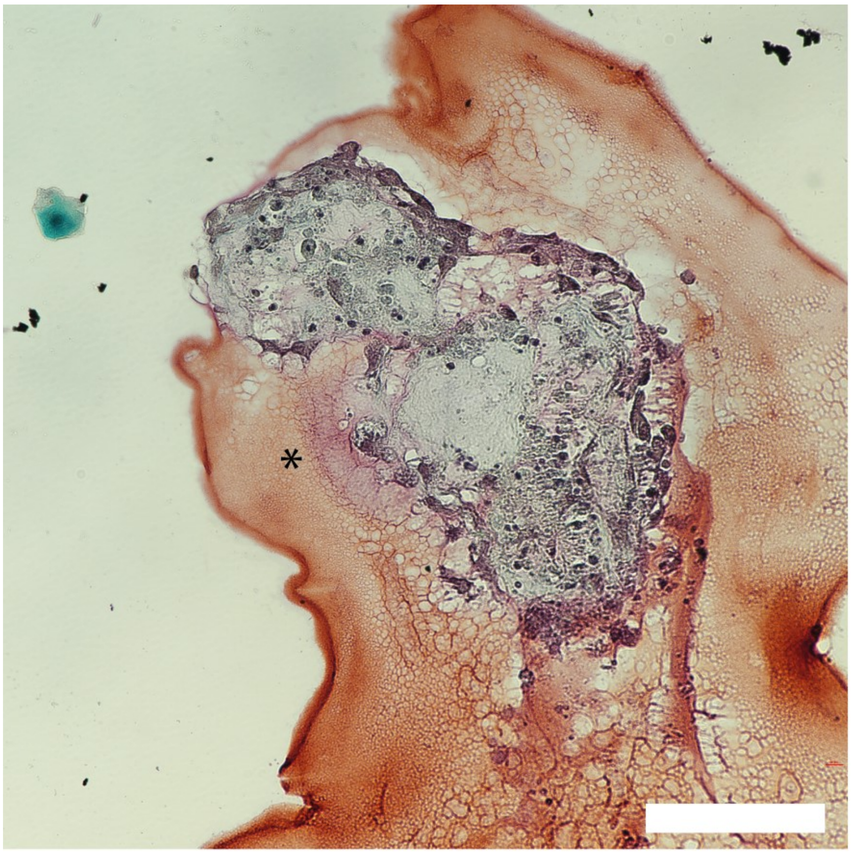


**Figure S3.** Enlarged view of Safranin O (SAF O) staining of HAMA-encapsulated hPDC spheroids on day 21. Image shown at the same magnification as the main figure with SAF O-positive regions highlighted with an asterisk. Scale bar is 100 µm.

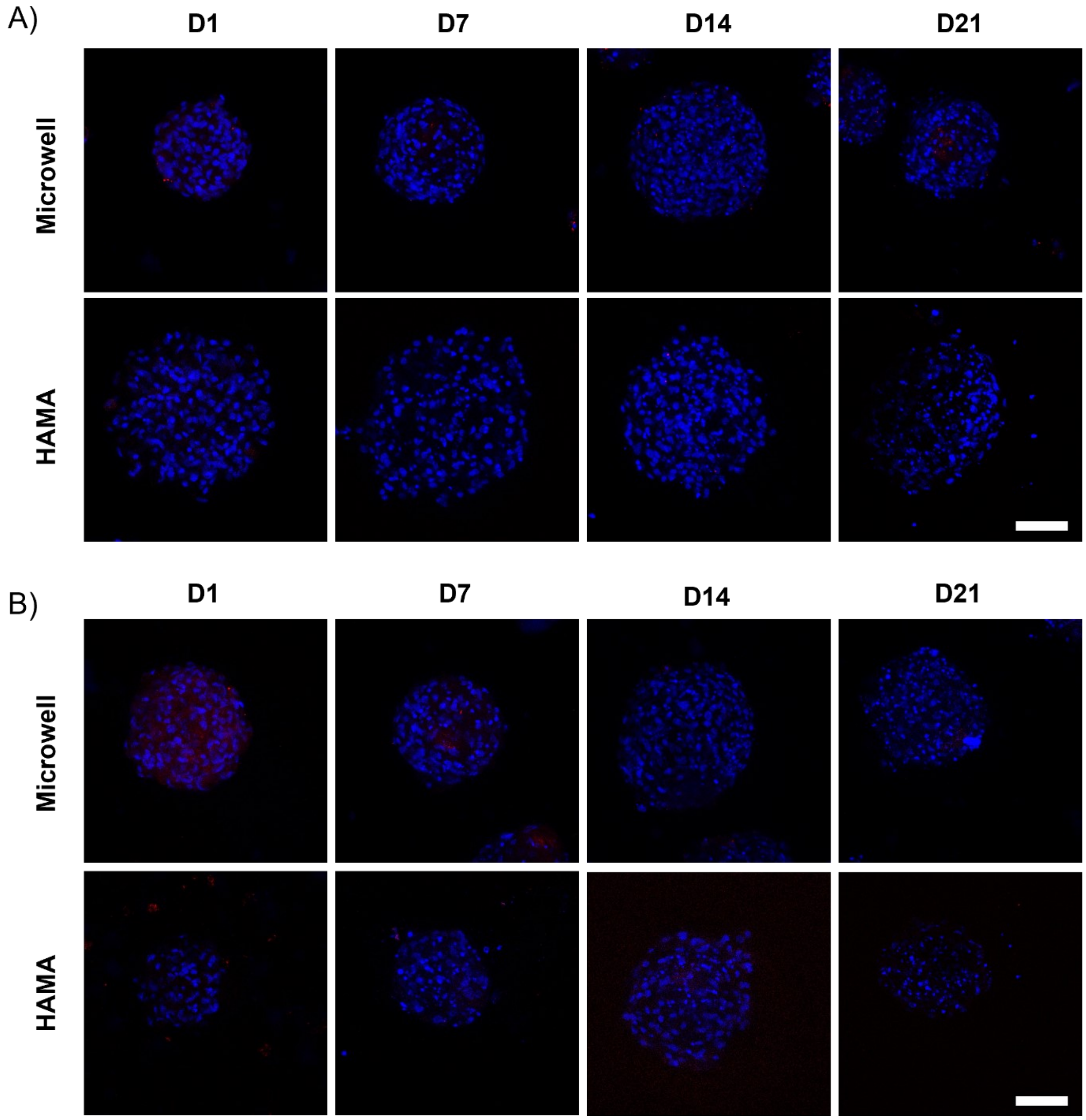


**Figure S4.** Representative maximum projection images of z-stacks obtained by confocal immunofluorescence staining of negative controls for hPDC spheroids on days 1, 7, 14 and 21. Negative controls included samples where no primary antibody was added (A) or where an IgG isotype control antibody was used (B). Cell nuclei were stained with DAPI (blue). Scale bars are 100 μm.

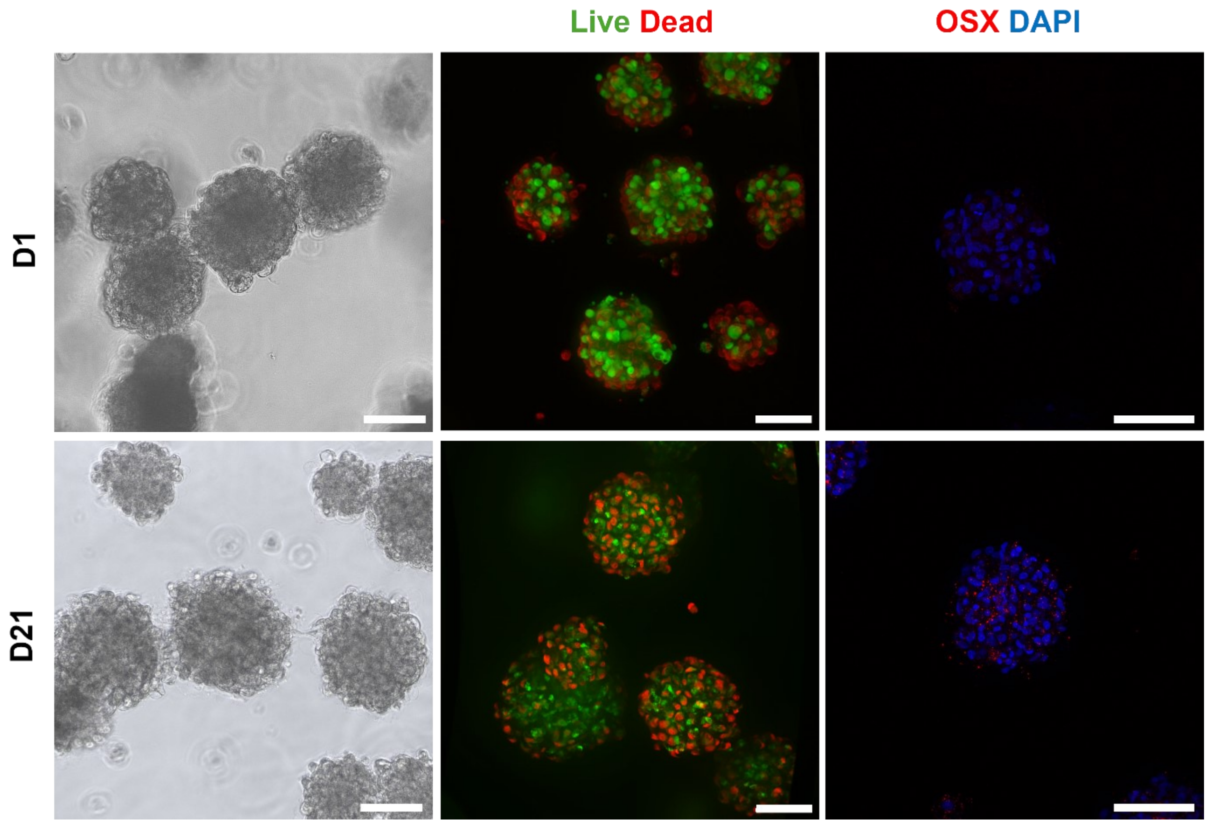


**Figure S5.** hPDC spheroids were bioprinted with a 250 µm nozzle. Bright-field images on days 1 and 21 (left), and live-dead staining on days 1 and 21 (middle). Live cells were stained with Calcein-AM (green), and dead cells were stained with EthD-1 (red). Representative maximum projection images of z-stacks obtained by confocal immunofluorescence staining for Osterix (OSX, red) on days 1 and 21 (right). Cell nuclei were stained with DAPI (blue). Scale bars are 100 µm.

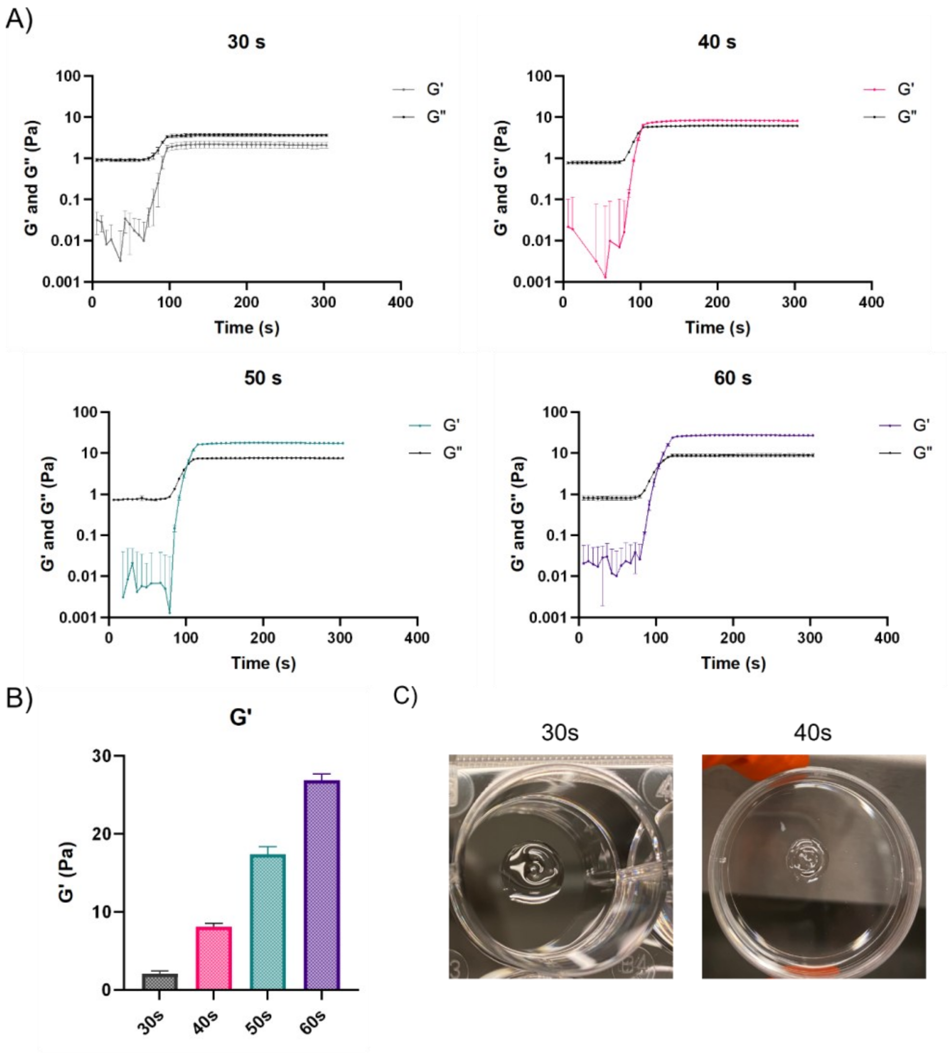


**Figure S6.** Photo-rheological characterization and bioprinting trials of partially crosslinked HAMA bioink. (A) Storage modulus (*G′*) and loss modulus (*G″*) of bioink containing 2% w/v HAMA and 0.01% LAP w/v in PBS crosslinked for 30, 40, 50, or 60 seconds. (B) Quantification of plateau *G'* for each crosslinking condition, presented as a column graph showing mean ± SEM (*n* = 3). (C) Bioprinting trial using bioink pre-crosslinked for 30 or 40 seconds.

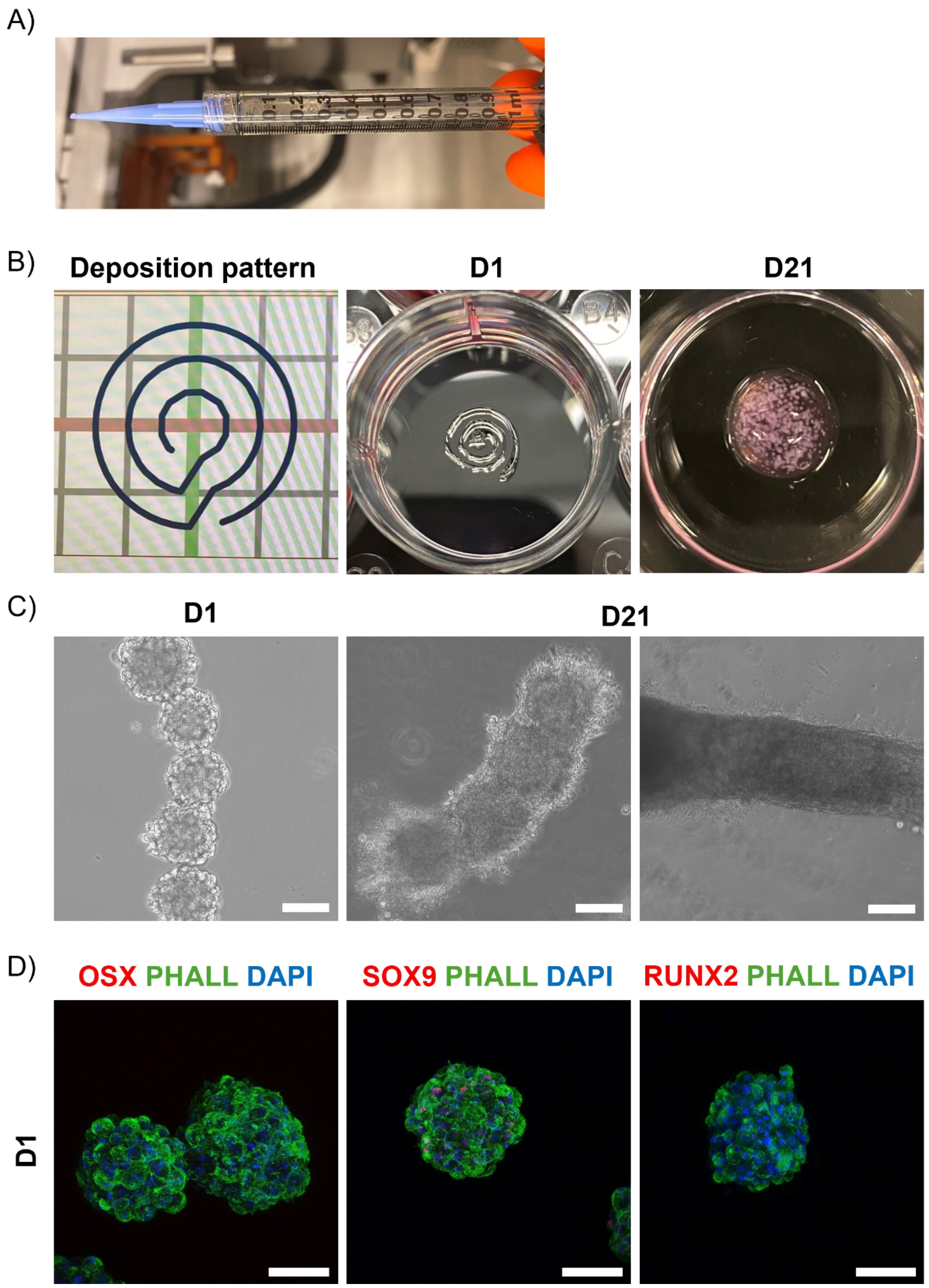


**Figure S7.** Macroscopic and microscopic analysis of bioprinted spheroid-laden HAMA constructs. (A) Syringe containing spheroid-laden bioink before bioprinting. (B) Deposition pattern used for bioprinting, shown as a set of connected circles with a 1 mm infill distance (left). The same pattern is visible in a one-layer bioprinted construct with spheroids on day 1 (middle). A four-layer bioprinted construct on day 21 (right). (C) Bright-field images of bioprinted constructs on day 1 and day 21, showing aligned spheroids on day 1 and partial or full fusion by day 21. (D) Representative maximum projection images of z-stacks obtained by confocal immunofluorescence staining at day 1 post-printing, showing DAPI (blue) for nuclei, phalloidin (green) for F-actin, and Osterix (OSX, red), SOX9 (red), or RUNX2 (red).

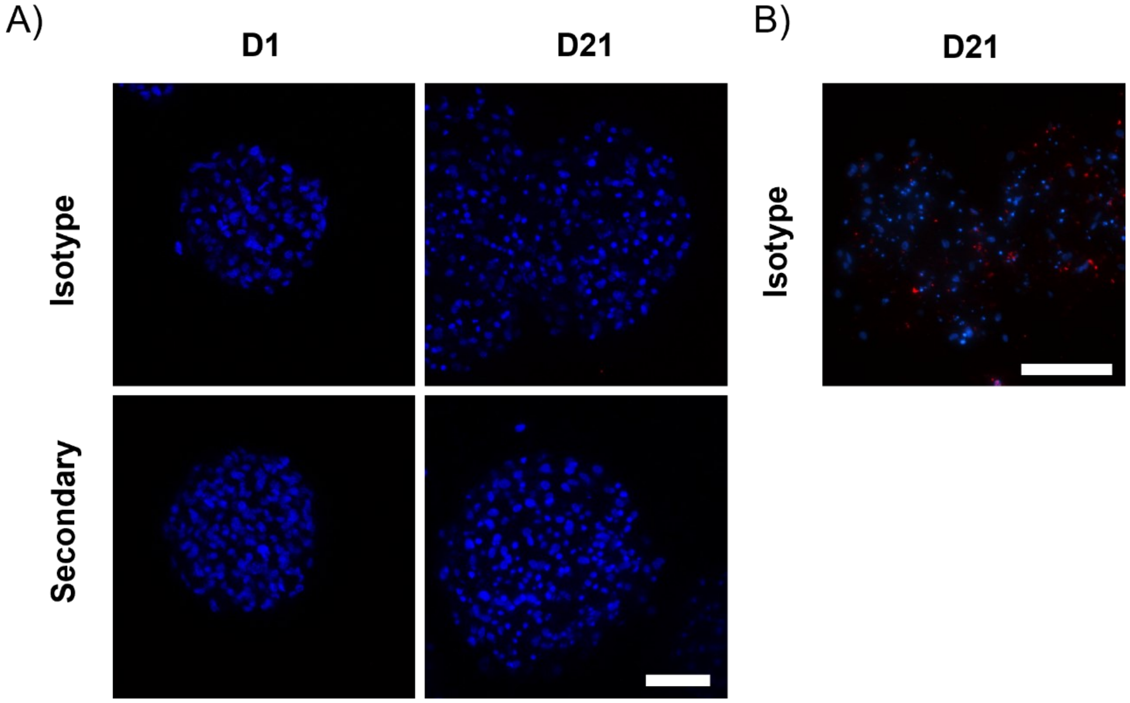


**Figure S8.** (A) Representative maximum projection images of z-stacks obtained by confocal immunofluorescence staining of negative controls for bioprinted hPDC spheroids on days 1 and 21. Negative controls included samples where no primary antibody was added or where an IgG isotype control antibody was used. (B) Representative immunostaining images of sectioned bioprinted hPDC spheroids stained with an IgG isotype control antibody. Cell nuclei were stained with DAPI (blue). Scale bars are 100 µm.
